# Bi-directional highways and super-seeder tissues underpin parasite dissemination in experimental visceral leishmaniasis

**DOI:** 10.64898/2025.12.30.696984

**Authors:** Ciara Loughrey, Juliana B T Carnielli, Shoumit Dey, Helen Ashwin, Sally James, Lesley Gilbert, Samantha Donninger, Alastair Droop, Grant Calder, Karen Hogg, Jon Pitchford, Jeremy C. Mottram, Paul M. Kaye

**Author notes:** **Correspondence:** Paul Kaye or Jeremy Mottram.

## Abstract

Visceral leishmaniasis (VL) is a life-threatening parasitic disease caused by *Leishmania donovani* and *L. infantum*. Although VL is characterised by organ-specific immunopathology and parasitism of the spleen, liver and bone marrow, other tissues may harbour parasites without overt pathology. The mechanisms governing patterns of within-host dissemination and tissue tropism are largely unknown. We used a barcoded library of isogenic *L. donovani* and an ecological analysis framework to evaluate parasite population diversity across tissues and to map parasite dissemination in a murine infection model. We reveal an unexpected high degree of inter-connectivity between parasite populations demonstrating: i) continual bi-directional dissemination between viscera and skin; ii) the existence of “super-seeder” sites that act as hubs fuelling systemic spread; and iii) rerouting of dissemination “highways” following immune perturbation. These findings change our understanding of *L. donovani* pathogenesis, providing critical insights into infection dynamics with direct implications for relapse, treatment failure and transmission.

## Introduction

The leishmaniases are vector-borne neglected tropical diseases, caused by intracellular protozoan parasites of the genus *Leishmania*. In mammalian hosts, these parasites primarily reside within myeloid cells, notably macrophages, monocytes, dendritic cells and neutrophils^1–7^, although evidence indicates they can also infect other cell types including haematopoietic stem cells (HSCs) and non-phagocytic cells such as fibroblasts^8–11^. Visceral leishmaniasis (VL), caused by *L. donovani* and *L. infantum*, is a life-threatening systemic form of leishmaniasis and is endemic in approximately 60 countries. Clinically, VL is characterised by gross pathology and high parasite loads in the liver, spleen and bone marrow^12,13^. Studies in experimental models^14–18^, reservoir species^19^ and patients, especially those immunocompromised by HIV co-infection^20–22^, indicate that parasites can disseminate widely even in the absence of overt local tissue pathology. The skin is particularly relevant, acting as an alternate route to blood for transmission^17,23^ and as the target tissue for post kala-azar dermal leishmaniasis^24–26^ (PKDL), a post-treatment sequela observed in 50-60% of VL patients in Sudan and up to 10% in India^26^. Beyond PKDL, the concept of persistent parasites residing in “sanctuary tissues” has also been suggested as a mechanism underpinning VL relapse and transmission, even after drug cure^9,27–29^.

Despite its clinical importance, little is known about within-host pathways of parasite dissemination into diverse tissues. Ecological and population dynamics principles offer a powerful framework for in vivo investigations to address these questions^30^. Recent advances in CRISPR genome editing have enabled genetic “barcoding” and the creation of isogenic libraries of pathogens with neutral, distinguishable alleles. This technique allows for detailed tracking of pathogen population dynamics in experimental models^31^, including through imaging^32–34^, lineage tracing^35^, experimental evolution^36^ and competition-based screening^37–42^. Sequence-tag based analysis of microbial population dynamics (STAMP) was initially developed by Abel et al^43^ and was described in the context of bottleneck quantification in a *Vibrio cholerae* rabbit infection model; the analytical approach involved the adaptation of previous work by Cavalli-Sforza and Edwards^44^ and Krimbas and Tsakas^45^ to analyse colonisation bottlenecks and relationships between pathogen populations in different tissue “niches”. The approach has since been used with other bacterial pathogens including *Pseudomonas aeruginosa*^46,47^, *Citrobacter rodentium*^48^, *Listeria monocytogenes*^49–51^*, Salmonella enterica*^52^*, Klebsiella pneumoniae*^53^*, Yersinia pseudotuberculosis*^54^ and *Escherichia coli*^55^, but to date has only been applied to one eukaryotic pathogen, *Toxoplasma gondii*^56^.

Here we leveraged this approach by generating an isogenic barcoded library of *L. donovani* to map systemic dissemination in a C57BL/6J murine experimental model of VL. We developed a network-based analytical approach to infer parasite population spread between tissues. Our findings reveal an unexpected degree of bi-directional communication between the skin and visceral organs and identify specific “super-seeder” tissues which act as major sources for widespread dissemination within a host. Furthermore, using a re-infection model we demonstrate that dissemination routes can be significantly perturbed by secondary exposure. These data provide fundamental insights into *L. donovani* pathogenesis, offering potential explanations for observed treatment failure and disease relapse.

## Results

### Generation of an isogenic, barcoded L. donovani library

To track parasite dissemination in vivo, we engineered an isogenic barcoded library of *L. donovani* promastigotes using CRISPR-Cas9 genome editing. We inserted 20bp barcodes into a neutral exogenous locus (**Figure 1a**). To determine the necessary library size for accurate Founder Population (FP) estimation using STAMP, we combined prior knowledge of the limitations of macro-ecological analysis (predominantly designed for relatively small numbers of “species”)^57^ and previous similar studies^49–51,56^ with a MATLAB-based simulation (see Methods for details). These simulations modelled various library sizes (10-500 barcodes) and scenarios of even and varied growth of individual parasites (**Figure 1b)**. We modelled the impact of random clonal expansion by selecting either 1 in 100 (common), 1 in 10000 (medium) or 1 in 1000000 (rare) individual parasites post-bottleneck to replicate at an enhanced rate of ten times the standard. Except in cases of common clonal expansion (which we anticipate is unlikely to occur in amastigote *Leishmania*), we observed that the accuracy of FP estimates was consistently high for all library sizes up to estimated FPs of 5000 (**Figure 1c** and **Supplementary Tables 1a and 1b**; Kendall correlation for FP values below 5000: all tau values above 0.83, p < 0.0001 for simulations other than common clonal expansion). Whilst FPs above this value cannot be discriminated effectively without a larger library, FPs below this point can be quantified using libraries within the limits tested here. We generated a library with 102 unique barcodes, with successful integration validated by PCR, and for a selected few, Sanger sequencing (**Figure 1a** and **Extended Data Figure 1a**). We confirmed that the inserted barcodes did not induce changes in parasite fitness. Culturing the full library for 7 days showed no evidence of any individual barcodes exhibiting a trend to increase or decrease beyond what would be expected to occur at random (**Figure 1d**). Furthermore, in vitro infectivity assays using murine bone marrow derived macrophages (BMDMs) demonstrated that all lines assessed (n=5) had similar distributions of parasites per macrophage (**Figure 1e** and **Figure 1f** and **Extended Data Figure 1b;** lambda parameters within 2 standard deviations at 1- and 3-hours post-infection), confirming the neutrality of barcoded lines for in vivo studies.

**Figure 1:**
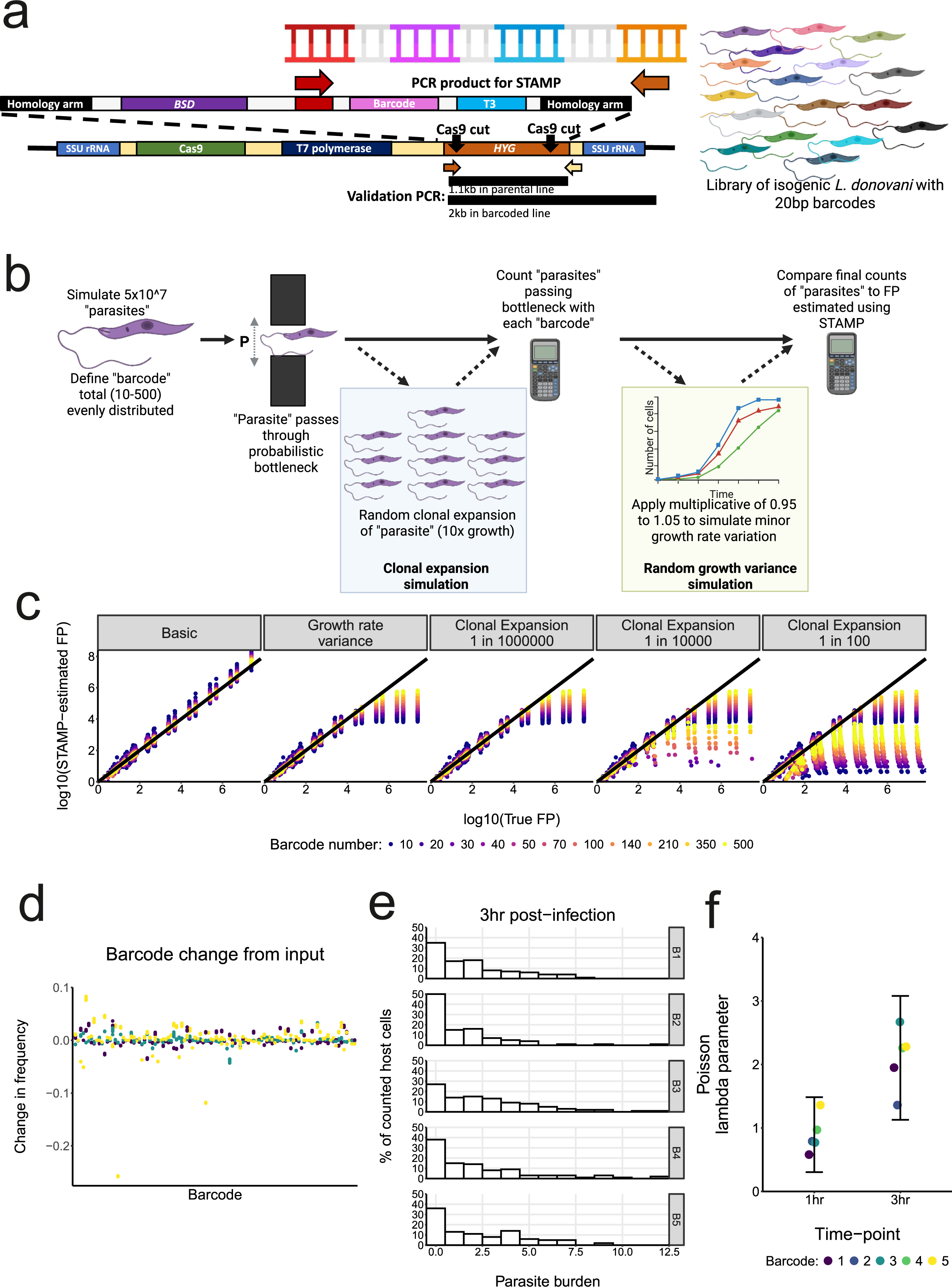
Use of CRISPR-cas9 genome editing to create a library of neutrally barcoded *L. donovani* promastigotes **a,** Schematic representation of the barcode insertion strategy for *L. donovani*. 20bp barcodes were inserted alongside a blasticidin resistance gene (BSD), by replacing the hygromycin resistance gene within the T7 cas9 cassette of the parental line. Coloured promastigote parasites indicate different barcodes in the library pool. A size-based validation PCR was used to validate successful integration of the barcode cassette into the desired location. Primer binding sites for this PCR are shown in orange and beige to match their respective binding sites. Primers for amplification of the barcode sequence for subsequent Illumina sequencing and Bar-seq analysis are shown in red and orange, to match to their respective binding sites. **b,** A schematic overview of the simulation strategy for assessing optimal library size. Simulations tested Founder Population (FP) calculation accuracy with: an even replication rate post-bottleneck; a 5% random variance in overall growth; and a simulation with 5% random variance plus random clonal expansion of 1 in 1000000 (rare), 1 in 10000 (medium) and 1 in 100 (common) individuals, with a clonal expansion growth rate simulated as 10 times greater than that of other individuals. **c,** STAMP-calculated estimates of FP against the true counted FP from the simulations in b. Colours indicate the number of barcodes in the library tested as shown in the key. Dark grey line indicates perfect accuracy of estimation (true FP is equal to STAMP-estimated FP). **d,** Frequency change from input frequency for each barcode in barcoded *L. donovani* library pools cultured as promastigotes for 7 days (n=5 for each input pool) minus total counts for each in the input culture pool. Colour indicates the experiment (three independent replicate experiments). For each barcode, each point indicates one of the five replicates within an experiment. **e,** Histograms of in vitro parasite infection burdens after a 3 hr incubation of promastigote parasites with murine bone marrow derived macrophages for five individual barcoded parasite lines, here termed B1-5. **f,** Poisson-fitted lambda values for the distribution of parasites per macrophage for each barcoded line infection for 1 or 3 hrs (as described in e). Colour indicates the individual barcoded parasite line in f. Error bars indicate 2x standard deviation for lambda values at each time-point.

### Tissue parasite population diversity reflects stage of infection

To examine temporal and spatial changes in parasite populations over the course of infection, we injected amastigotes of barcoded *L. donovani* intravenously into female C57BL/6J mice (**Figure 2a**), as this is an established model for generating reproducible systemic infections^58^. Our broad aim was to apply diversity measures both within and between tissue samples within an individual animal, and to use this information to perform network analysis on the “ecosystem” of parasites within the host (**Figure 2a**).

**Figure 2:**
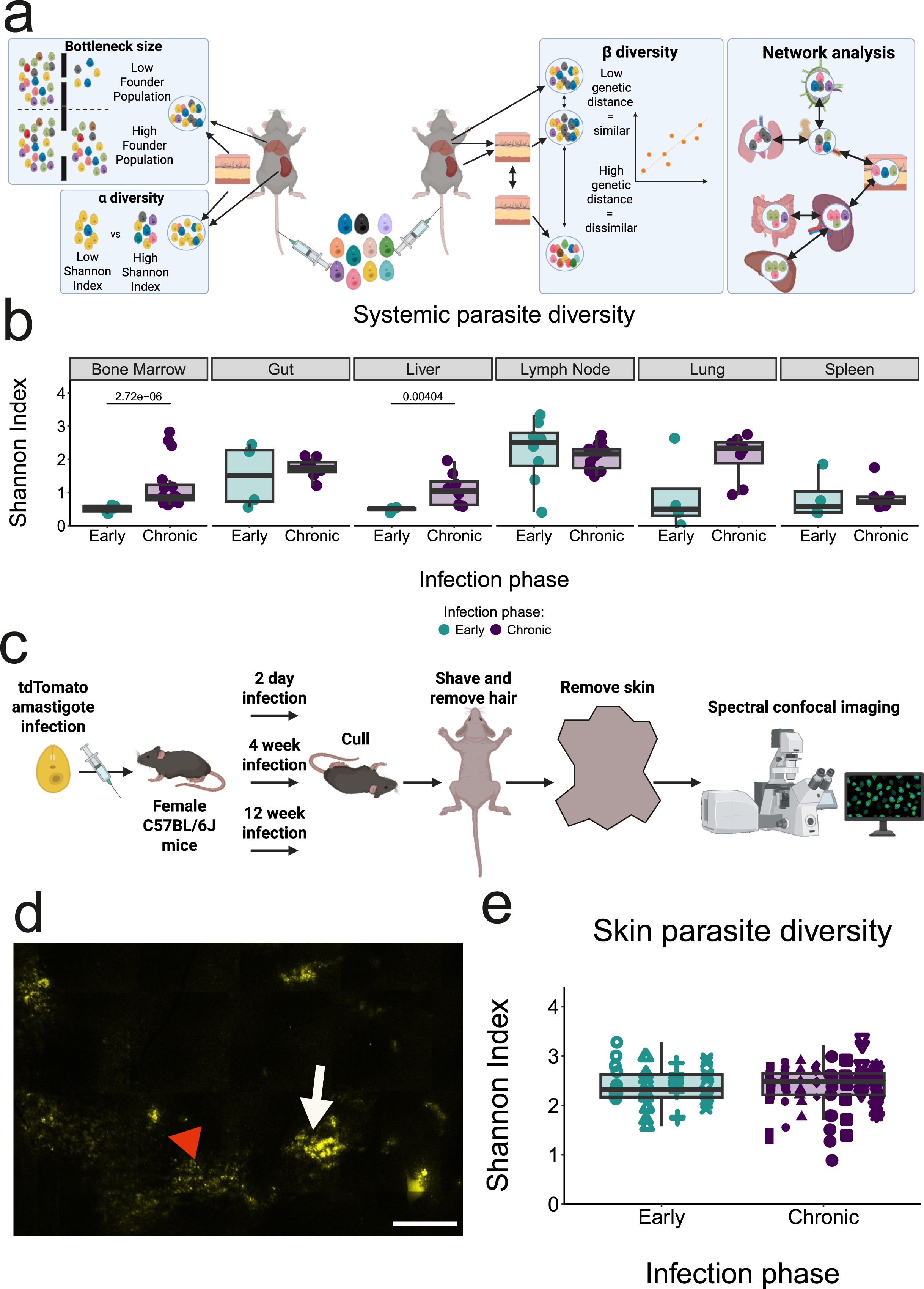
Parasite population structure varies both spatially and temporally **a,** Schematic illustrating macro-ecology and graph theory concepts applied in this study. **b,** Parasite diversity (Shannon Index [SI]) in systemic tissues of female C57BL/6J mice infected with barcoded amastigotes for 4 weeks (early; n=4; turquoise) or 10-12 weeks (chronic; n=8; purple). For each mouse, two lymph nodes and two bone marrow samples were analysed and as such n number is doubled for visualisation and statistical analysis. **c,** Schematic of protocol for analysis of skin parasite distribution. n=4 each for 2 day and 4 week infections. n=3 for 12 week infections. **d,** Representative image (after stitching of panels) of a section of late infected skin. Parasites are identified by high (white arrows) and low (red arrowheads) tdTomato signal (yellow). Scale bar represents 2000μm. **e,** Parasite diversity (SI) in skin biopsies (12 per mouse) for the mice described in panel b (early; n=4 and chronic; n=8); all 12 data points per mouse are shown and used for statistical comparison. Shapes indicate individual mice. All boxplots show the median, 25th and 75th quartiles, with whiskers representing all data-points within 1.5x IQR of the hinges; points above or below the whiskers represent outlying data values. p-values from a two-tailed Mann-Whitney-Wilcoxon test for within tissue time-point comparisons.

We first used the concept of alpha (within site) diversity^59–61^ to analyse parasite populations at 4 weeks (early) and 10-12 weeks (chronic) post-infection (p.i) (**Figure 2b**). Our analysis revealed a significant increase in parasite diversity (quantified using the Shannon Index, which measures both richness and evenness of the population^59,61^) in bone marrow (p < 0.0001) and liver (p = 0.00404) (**Figure 2b** and **Supplementary Table 2**). Early post-infection, spleen, liver and bone marrow parasite populations showed a relatively consistent degree of diversity between mice, whereas both median diversity and inter-mouse variation was greater in the lymph node and gut. This may reflect different kinetics or routes for colonisation in these tissues. These findings are important as they suggest that tissue colonisation is not governed by a single bottleneck event, but rather reflects a continuous process of recruitment and colonisation throughout the course of infection.

We have previously shown that parasites are located in “patches” in the skin of immuno-compromised *B6.Rag2^−/−^* mice^17,62^. We therefore adopted the same analytical approach to visualise skin parasites during early and chronic infection in immuno-competent C57BL/6J mice, using a tdTomato-expressing *L. donovani* line (**Figure 2c**). Confocal imaging detected parasite patches only during chronic infection (**Figure 2d**, **Extended Data Figure 2** and **Supplementary Figures 1 and 2**). Of note, heterogeneity in the tdTomato signal was observed. This may reflect differences in patch parasite burden and/or the presence of quiescent parasites with reduced metabolic activity^8,28,29^. To examine parasite population diversity in the skin, we sequenced 12 spatially adjacent biopsies per mouse taken from the flank. The average parasite diversity in these biopsies was similar between early and chronic infection. However, the diversity of biopsies derived from a single mouse was highly heterogeneous, with individual mouse being the best predictor of parasite diversity in skin biopsies, rather than time post-infection (**Figure 2e**; Compound Poisson Generalised Linear Mixed Model [GLMM], see **Supplementary Note**). Overall, the diversity in skin, lymph nodes, lung and gut was higher and more variable than in the bone marrow, spleen and liver (**Figure 2b**), supporting a model of stochastic and continuous seeding of the skin, gut and lymph nodes from visceral tissues^17,18,62^.

### Skin patches vary in their similarity to parasite populations in other tissues

We next used the ecological concept of beta (between site) diversity, calculated via genetic distance [GD] using the Cavalli-Sforza chord distance^44^) to quantify the relatedness between parasite populations in different tissues (**Figure 3a**). A heatmap representation confirmed extensive tissue inter-connectivity, though the specific pattern varied widely between individual mice. For example, in the representative mouse shown in **Figure 3b**, the liver parasite population is more highly related to that of the lung and spleen than to other tissues. Parasites in skin biopsy 03 are most similar to those in bone marrow and to a lesser extent other skin biopsies.

**Figure 3:**
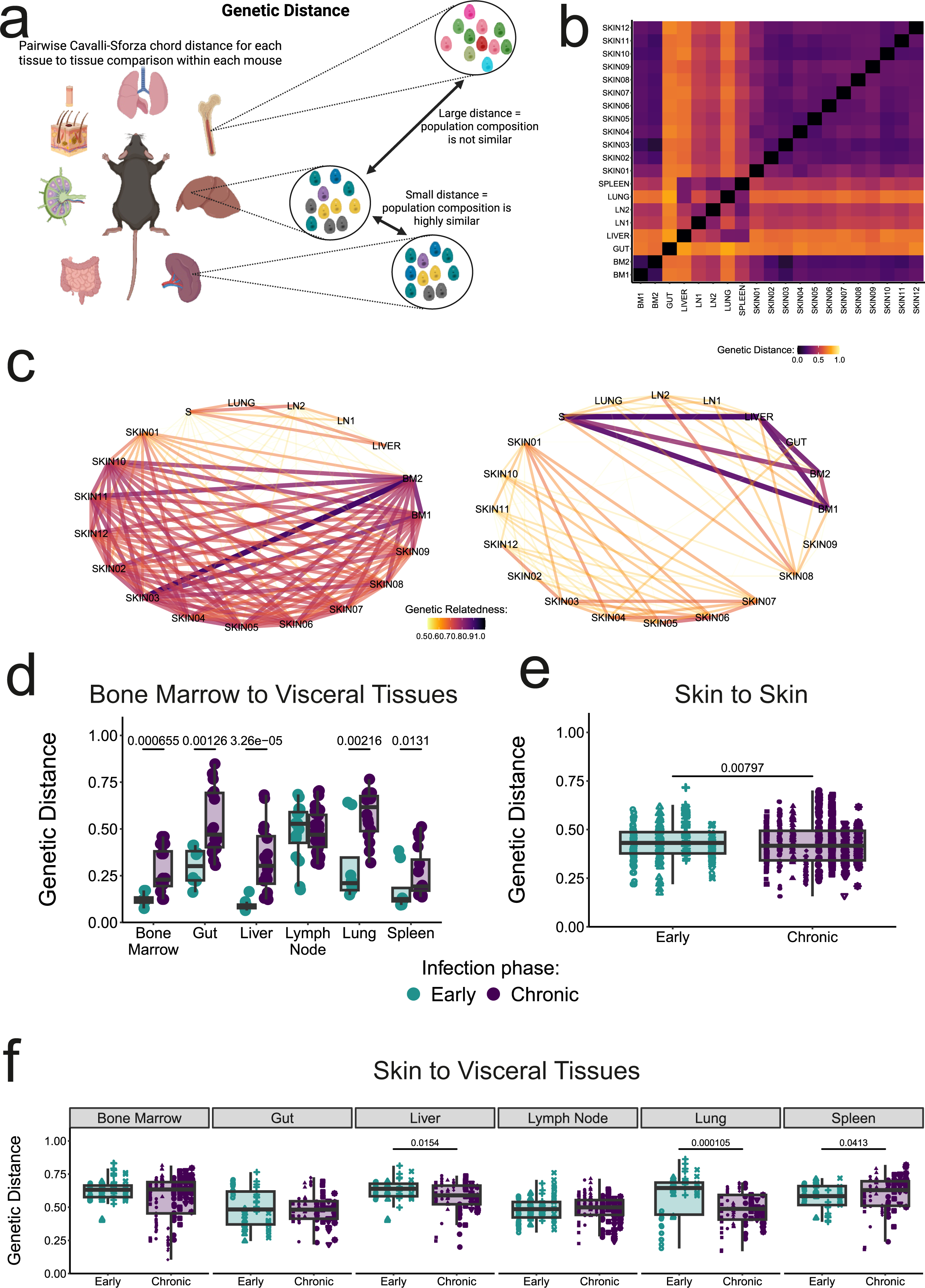
Genetic distance analysis reveals heterogenous skin landscape, with increasing connectedness over the course of infection **a,** Schematic of genetic distance (GD) concept for pairwise comparison of tissue parasite populations in individual mice. **b,** Example heatmap for a single mouse at 12 weeks post-infection. **c,** Network representations of GD (edges) between parasite populations in each tissue (vertices). Networks for two exemplar mice (Mouse 21 and Mouse 23) at 12 weeks post-infection, with all GD above 0.5 removed. Line colour indicates genetic relatedness (1-GD) as shown in the legends for each network. Thicker, more opaque connecting lines indicate a higher relatedness (lower GD). **d,** GD for bone marrow and all other systemic tissues in female mice with early (turquoise; n=4) or chronic (purple; n=8) infection. For each mouse, two lymph nodes and two bone marrow samples were analysed, meaning n number is doubled for visualisation and statistical analysis. **e,** GD for pairwise comparisons of each skin biopsy with all other skin biopsies within each individual mouse (n=12 biopsies per mouse). **f,** GD for pairwise comparisons of each skin biopsy with visceral tissues within each individual mouse (n=12 biopsies per mouse). Shape indicates individual mice. Colour legend applies to d, e and f. In b and c, BM = bone marrow, S = spleen and LN = lymph nodes. All boxplots show the median, 25th and 75th quartiles, with whiskers representing all data-points within 1.5x IQR of the hinges; points above or below the whiskers represent outlying data values. p-values from a two-tailed Mann-Whitney-Wilcoxon test for within tissue time-point comparisons. GD statistical testing: Bone Marrow to Bone Marrow: p = 0.000655; Bone Marrow to Gut: p = 0.00126; Bone Marrow to Liver: p < 0.0001; Bone Marrow to Lung: p = 0.00216; Bone Marrow to Spleen: p = 0.0131; Skin to Skin: p = 0.00797; Skin to Liver: p = 0.0154; Skin to Lung: p = 0.000105; Skin to Spleen: p = 0.0413.

To visualise the parasite “ecosystem” in individual mice, we used 1-GD (i.e. genetic relatedness) as a connection strength measure for a graphical network (**Figure 3c** and **Extended Data Figure 3**). We found that while visceral tissues were consistently linked during early infection (**Extended Data Figure 3**), the linkage between skin and viscera became highly heterogeneous during chronic infection (**Figure 3c** and **Extended Data Figure 3**). We further noted that bone marrow became more genetically distant from all other visceral tissues, aside from lymph nodes, over time (**Figure 3d**). With the exception of the gut becoming more distant from spleen and liver, no other visceral pairs showed significant changes in relatedness (**Supplementary Table 3**). We also found that the average relatedness of the parasites in each skin biopsy was greater (reduced GD) at chronic infection compared to early infection, but again high intra-mouse variation made individual a better predictor of inter-skin relatedness (**Figure 3e**, **Supplementary Table 3** and **Supplementary Note**).

A distinct clustering effect was observed when comparing skin biopsies to visceral organs; some mice had skin patches highly related to one key visceral tissue, whilst others showed low relatedness (**Figure 3f**). Some of these visceral tissue-skin relationships also changed over time when we compared individual biopsy-viscera GDs. We also noted tissue-specific differences had an interaction effect with time p.i (**Supplementary Table 3** and **Supplementary Note**). This suggests that the relationship between parasites in different visceral tissues and in skin varies over time in a manner that depends on the tissue in question. Furthermore, comparison of GD to the original input pool indicated that a single colonisation event from our original injection pool could not account for the patterns we had observed across tissues in our parasite population diversity and GD analysis (**Extended Data Figure 4**), indicating that later tissue-to-tissue dissemination does indeed play a role in parasite tissue colonisation patterns.

### Identification of ‘super-seeder’ sites

To infer directionality of parasite dispersal events within a host, we developed a new analytical approach (**Figure 4a** and Methods). Our metric Tissue-Sourced FP (TSFP) calculates the FP for a tissue with reference to an input population from another tissue and can be used in a pairwise manner to infer routes of parasite trafficking within each mouse. This approach relies on the overall divergences between barcode frequencies to calculate a probable direction and relative size of pathogen movement between tissues.

**Figure 4:**
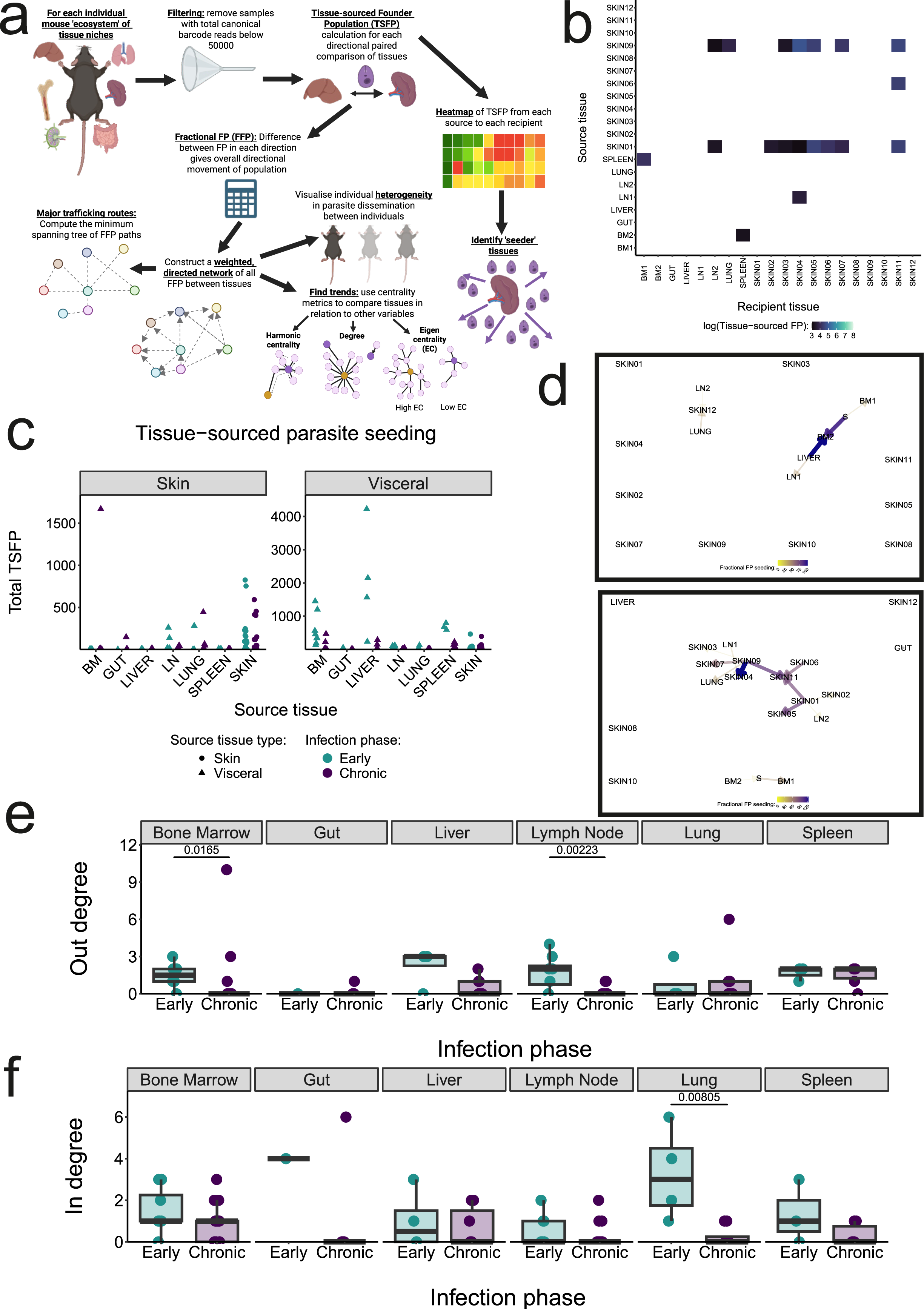
Fractional Founder Population analysis reveals ‘super-seeder’ parasite populations **a,** Schematic illustration of the analytical pipeline to visualise and assess parasite dissemination pathways. **b,** Exemplar heatmap (Mouse 24) for TSFP for an individual mouse at 12 weeks post-infection. Scale represents log(TSFP) after thresholding applied (cut-off TSFP = 20) for each directional tissue to tissue relationship. White indicates those TSFP values below threshold or tissues which were excluded in filtering. **c,** Total TSFP sourced from each tissue sample to either skin (left) or visceral (right) recipient tissues. Data from early (turquoise; n=4) or chronic (purple; n=8) infection are shown. **d,** Minimum spanning tree (MST) network representations of FFP for two exemplar mice (Mouse 22 and Mouse 24) at 12 weeks post-infection. Legends, line colour and thickness indicate FFP. Arrowheads represent direction of FFP. **e, f,** Out-degree (e) and in-degree (f) values for each visceral tissue in the full FFP networks constructed for each mouse. infection phase colour legend applies to c, e and f. In b, c and d, BM = bone marrow, S = spleen and LN = lymph nodes. All boxplots show the median, 25th and 75th quartiles, with whiskers representing all data-points within 1.5x IQR of the hinges; points above or below the whiskers represent outlying data values. All statistics shown are p-values from a two-tailed Mann-Whitney-Wilcoxon test for within tissue comparisons between time-points.

Heatmap visualisation of TSFP values (**Figure 4b** and **Extended Data Figure 5**) revealed that many mice exhibited one or two ‘super-seeder’ sites, acting as major parasite sources for other sites. For example, in the mouse shown in **Figure 4b**, skin biopsies 01 and 09 act as a source for parasites found in multiple other skin biopsies, as well as lung (biopsy 09) and lymph node (both biopsies). While the liver and bone marrow predominantly acted as sources for parasites in other visceral tissues, both visceral organs and some skin sites could act as super-seeders. Importantly, the skin was found to seed both other skin sites and, critically, visceral tissues (**Figure 4c**). In relation to our simulation findings showing a lack of discrimination power above 5000 where growth rate is variable, we noted that all calculated TSFPs were below this level and thus not impacted by this confounder.

We then calculated the difference between the TSFP values in each direction for each pair of tissues; we termed this metric Fractional Founder Population (FFP). Using FFP to build directed networks, we visualised the major “highways” of parasite dissemination (**Extended Data Figure 6**) via a minimum spanning tree (MST) analysis (**Figure 4d** and **Extended Data Figure 7**). This revealed heterogeneity in dissemination patterns between individual mice. For example, as shown in **Figure 4d**, the parasite “highway” may extend along a visceral axis, or be predominantly within skin. Using the complete FFP networks (**Extended Data Figure 6**), we next calculated for each visceral tissue the “out-degree” (i.e. the number of outward connections, signifying a tissue’s ability to act as a parasite source) and “in-degree” (i.e. the number of inward connections, signifying receipt of parasites from elsewhere). Supporting early observations that suggested increasing isolation of the bone marrow (**Figure 3d**), this analysis showed that out-degree decreased significantly during chronic infection (p = 0.0165; **Figure 4e**; **Supplementary Tables 4** and **5**). We also found that the lymph nodes and lung exhibited a reduction in out- and in-degree respectively (p = 0.00223 and p = 0.00805; **Figure 4e** and **Figure 4f**; **Supplementary Tables 4** and **5**). Whilst not statistically significant, we noted a trend towards reduced out-degree in the liver, which is likely a reflection of its transition to control of parasite burden in chronic infection.

### Skin parasites may act as a ‘re-seeder’ of visceral organs

Our prior GD analysis (**Figure 3f, Supplementary Table 3** and **Supplementary Note**) indicated that skin parasite populations may be interacting with visceral tissues in a dynamic manner. Further TSFP analysis confirmed dynamic, bi-directional communication between the skin and visceral organs (**Figure 5a** and **Figure 5b**). For viscera to skin seeding, this revealed a characteristic pattern where skin patches were seeded from one or two key visceral tissues, with the source tissue varying between individual mice (**Figure 5a**). However, comparing total TSFP seeding to the sum of all TSFP to skin biopsies for each mouse indicated no significant change between early and chronic stages for any specific source tissue (**Supplementary Table 6**). Total skin-to-spleen and skin-to-lung seeding decreased during chronic infection (**Figure 5b** and **Supplementary Table 7**). Skin-to-skin seeding across all biopsies was greater during chronic infection (p = 0.00257; **Supplementary Table 8**), but there was no significant difference between early and chronic infection when comparing total skin seeding (**Figure 5c** and **Supplementary Table 9**). This indicates that localised parasite spreading is driven by a greater number of “super-seeder” biopsies, rather than an evenly distributed increase in seeding across all biopsies.

**Figure 5:**
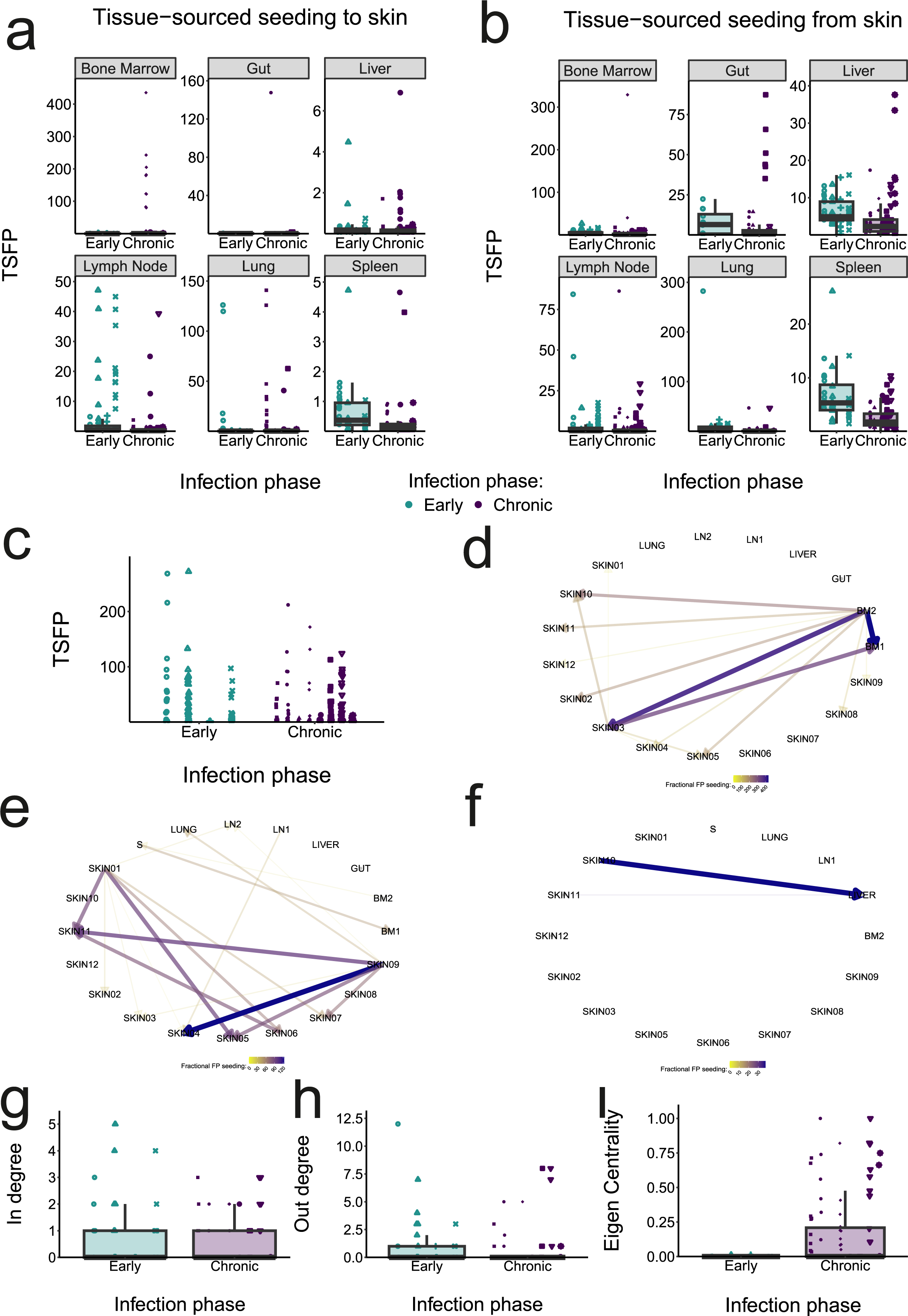
Evidence of skin parasite spread within the skin and to visceral organs **a, b,** TSFP from visceral tissues to skin (a) and from skin to visceral tissues (b) for early (turquoise; n=4) or chronic (purple; n=8) mice. **c,** Skin-to-skin seeding, quantified as TSFP, taken from each mouse described in a and b (n=12 biopsies per mouse). **d, e, f,** Full networks constructed using FFP as the metric for edge weight, for Mouse 21 (d), Mouse 24 (e) and Mouse 25 (f). Line colour indicates FFP as in legends. Thicker, more opaque lines indicate a higher FFP in the direction indicated by the arrowhead. **g, h, i,** In-degree (g), out-degree (h) and eigen centrality (i) for skin sites in the FFP networks. Shape indicates biopsies from each individual mouse within each time-point group. Infection phase colour legend applies to a, b, c, g, h and i. In d, e and f, BM = bone marrow, S= spleen and LN = lymph nodes. All boxplots show the median, 25th and 75th quartiles, with whiskers representing all data-points within 1.5x IQR of the hinges; points above or below the whiskers represent outlying data values.

Network construction revealed that the skin-viscera interaction axis differs strikingly between individual mice (**Figures 5d-f** and **Extended Data Figure 8**). For example, in one mouse the bone marrow is acting as a parasite source for multiple skin biopsies (**Figure 5d**). In another a single skin biopsy is the dominant source for other skin sites (**Figure 5e**) and a third illustrates the capacity of skin to seed the liver (**Figure 5f**). Analysis of in-degree and out-degree found no change between early and chronic infection for skin biopsies (**Figure 5g; Supplementary Table 5**; **Figure 5h; Supplementary Table 4**). Comparing eigen centrality, a measure of connectedness to other highly connected vertices in a network (see **Figure 4a**), we saw a clear increase in the number of highly connected sites in chronic infection, although the average value across skin biopsies was not significantly different (**Figure 5i; Supplementary Table 10**). This suggests that specific skin sites become more closely connected to the core dissemination network over time, despite exhibiting no change in average connection numbers.

### Immune perturbation via secondary infection alters skin seeding dynamics

To explore the stability of these dissemination patterns, we investigated how they were affected by immune perturbation resulting from secondary infection. Mice infected for 10 weeks with barcoded amastigotes were re-infected with luciferase-expressing *L. donovani*. Uninfected mice were used as a control for infectivity of the luciferase-expressing parasites and to assess the degree of protective immunity (**Figure 6a** and **Supplementary Table 11**). As expected, at two weeks p.i, ex vivo imaging indicated significant protection against secondary infection (**Figure 6b**, **Figure 6c**, **Extended Data Figure 10** and **Supplementary Table 11**).

**Figure 6:**
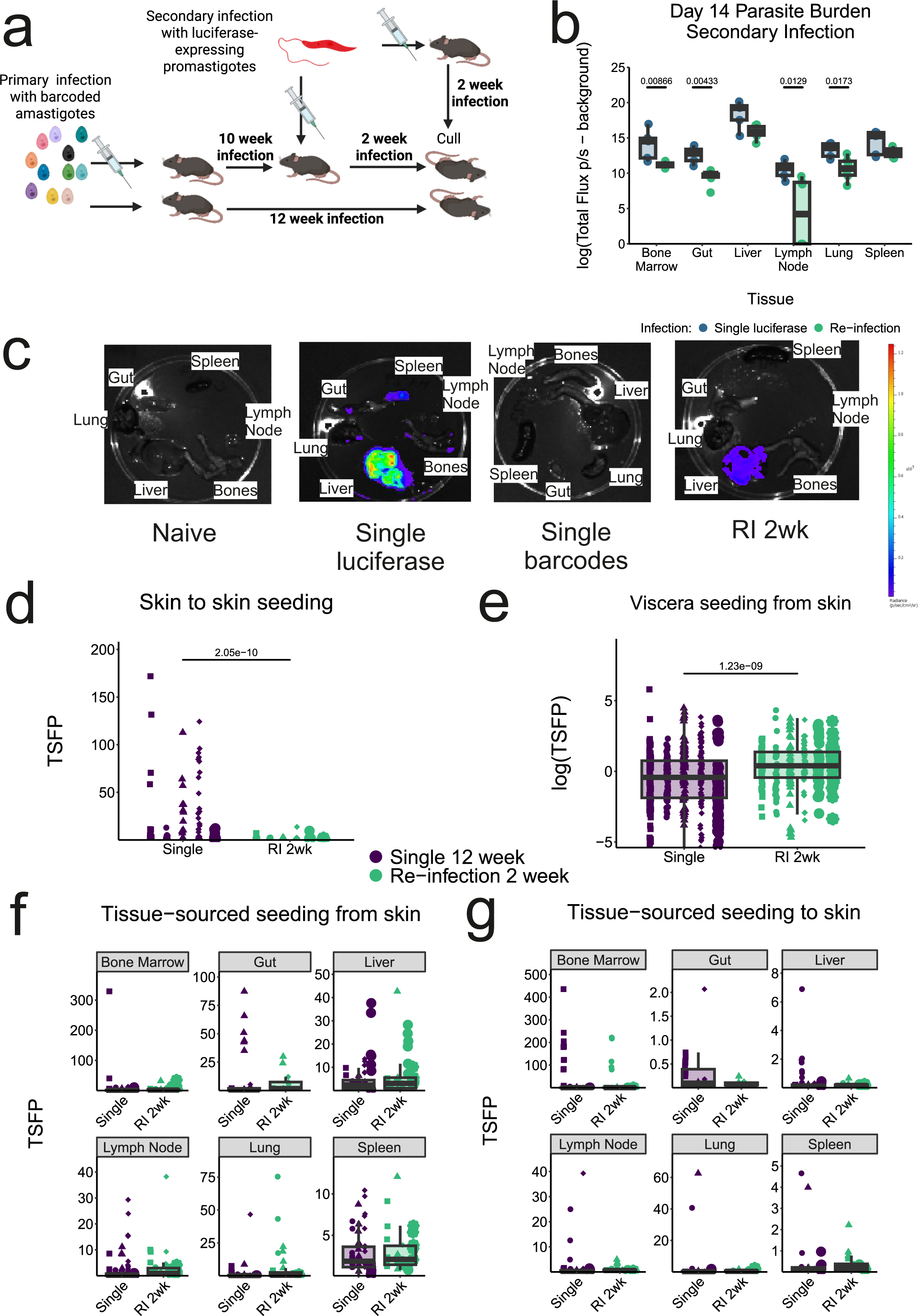
Secondary infection alters skin seeding dynamics to both local skin sites and visceral tissues **a,** Schematic of experimental design for re-infection of female C57BL/6J mice. Single 12 week infection: n=5 mice (also used as part of the previously described “chronic” infection group); 12 skin biopsies per mouse. Single 2 week infection: n= 5 mice. Re-infection (RI): n= 6 mice; 6 skin biopsies per mouse. **b,** Luciferase signal (total flux p/s) minus naïve tissue background signal for visceral tissues for mice after 2 week infection with luciferase-expressing *L. donovani* (single infection; blue) or the same infection after prior infection with barcoded amastigote *L. donovani (*re-infection; green). **c,** Representative ex vivo IVIS images of visceral tissues post-cull. **d,** TSFP for skin-to-skin seeding for each biopsy in the single (barcode only; purple) and re-infected (barcodes then luciferase; green) mice. **e,** log(TSFP) for skin-to-visceral tissue seeding (amalgamated for all visceral tissues) for each skin biopsy in the single and re-infected mice. **f, g,** TSFP for skin-to-visceral tissue seeding split by tissue recipient (f) and for visceral-to-skin tissue seeding split by tissue source (g) for each individual biopsy. Shape indicates individual mouse. The same colour legend applies to d, e, f and g. All boxplots show the median, 25th and 75th quartiles, with whiskers representing all data-points within 1.5x IQR of the hinges; points above or below the whiskers represent outlying data values. p-values from a two-sided Mann-Whitney-Wilcoxon test for total flux minus background (b) and for all TSFP skin biopsy pairs (d) and skin-viscera pairs (e) between the single (two week luciferase) infection group and the re-infection group (b) and for the single (12 week barcode) infection group and the re-infection group (d and e). IVIS total flux statistical testing (b): Bone Marrow p = 0.00866; Gut p = 0.00433; Lymph Nodes p = 0.0129; Lung p = 0.0173.

Re-infection had notable effects on the primary barcoded parasite population. First, inter-skin seeding within a patch was significantly lower after re-challenge (**Figure 6d** and **Supplementary Table 12**; MW test for TSFP of all biopsy pairs within each individual p < 0.0001), driven by a reduction in high-TSFP “super-seeder” events. Similarly, overall seeding decreased (total TSFP within a skin patch in single vs re-infection: p = 0.00433; **Supplementary Table 13**). This could indicate a reduced skin parasite load or alternatively, a reduction only in parasite spread within the skin. Second, overall skin to visceral tissue trafficking was significantly increased after re-infection (MW test for TSFP of all biopsy-visceral tissue pairs within each individual p < 0.0001; **Figure 6e**; **Supplementary Table 14**), suggesting the perturbed immune environment promoted parasite dissemination from the skin. When we compared total TSFP for each tissue from all skin biopsies in each mouse, we noted that bone marrow to skin seeding decreased after re-infection (p = 0.00896; **Supplementary Table 15** and **Supplementary Table 16**). Third, the impact of re-challenge on migration was tissue-specific (compound Poisson GLMM; **Figure 6f** and **Figure 6g**), whilst mouse-to-mouse heterogeneity was also a significant random effect (**Supplementary Note**). Collectively, these findings demonstrate a complex and extensive rerouting of dissemination pathways in response to immune perturbation. The skin appears to act as a dynamic parasite “sanctuary”, capable of re-seeding other tissues in specific immunological conditions, which has important implications for disease relapse and transmission.

## Discussion

In this study, we have employed CRISPR genome editing to generate a neutral barcoded *L. donovani* library and combined this with a novel network-based analytical approach to examine the dissemination patterns of *L. donovani* in a murine model of VL. This network analysis, which extends previous STAMP^43^ methodology, allowed us to infer the directionality of parasite migration within the host, revealing a highly inter-related dissemination network connecting parasite populations across different tissues.

Our findings highlight three previously unrecognised features of the host-*Leishmania* interaction, which change the understanding of parasite persistence, relapse and transmission. First, our network analysis demonstrates that the wide distribution of parasites across tissues, often viewed as a static consequence of systemic disease, is in fact merely a snapshot of a highly dynamic and inter-connected dissemination network. Although previous imaging studies using serial intra-vital imaging of *Leishmania*^63^ and *Trypanosoma cruzi*^64^ infections hinted at dynamic tissue tropism, our data formally establishes the inter-relationships between parasite population structures at distinct tissue sites. Our analytical approach allowed us to identify, on an individual host basis, common dissemination “highways” and notably revealed bi-directional seeding of parasites both within and between visceral organs and skin. Surprisingly, within this highly inter-connected network, the bone marrow parasite population becomes increasingly isolated from populations in other visceral organs as the infection progresses. This finding aligns with reports of high parasitism of haematopoietic stem cells in experimental VL^8,9^ and the concept that the bone marrow serves as a protective sanctuary for other persistent pathogens, such as *Mycobacterium tuberculosis*^65^ and *Plasmodium* parasites^66^, suggesting a mechanism for long term sequestration. A greater understanding of the mechanics underlying parasite persistence and inter-organ seeding would, if translatable to humans, provide potential treatment targets for parasite elimination in affected patients, and could be of particular benefit to those most at risk of disease relapse.

Second, we observed frequent seeding of parasites to the skin, a finding consistent with reports in immunodeficient mice^17,62^, dogs^67^ and humans^68^. Extending our previous work identifying a highly heterogeneous “patchy” distribution of parasites in the skin^17,62^, we now show that parasite populations sampled within distinct biopsies across the skin are interconnected and frequently linked to one or more visceral organs. These observations reinforce a model whereby skin parasite patches originate by initial stochastic seeding from the viscera, followed by localised patch spread, ultimately creating larger inter-related patches. Importantly, our analysis also identifies “super-seeder” sites, which act as disproportionate sources for seeding of multiple other tissues. Our analysis of multiple skin biopsies indicates that super-seeding may be restricted to one or a few biopsies per mouse. Given the intracellular lifestyle of *Leishmania* within mammalian hosts, we hypothesise that super-seeding behaviour likely reflects the downstream consequences of events triggering mass migration of myeloid cells, including those containing parasites, into the draining lymphatics and subsequently the circulation.

Experimental testing of this hypothesis remains challenging, given the current inability to track the fate of individual skin patches over time, or to manipulate myeloid cell migration in a site-specific manner. However, inflammatory monocytes offer a specific target cell for investigation, given their established role in experimental VL infections^2,69^. Previous studies have identified quiescent parasites in CL skin lesions^28,29^ and our imaging data is suggestive of parasite phenotypic heterogeneity across skin patches, opening the possibility that “super-seeder” sites may contain a more actively dividing and diverse parasite population ready for export. Identification of the cellular and molecular landscape that confers “super-seeder” status to some but not all sites is a major focus of current research, as is the role of “super-seeder” sites in facilitating transmission.

Although we extrapolate from rodents to humans with caution, it is also reasonable to suspect that similar principles may apply to dissemination across species barriers. Pre-clinical and clinical development of therapeutics for VL mandate efficacy against parasites in the major visceral tissues and treatment options are generally not selected based on their distribution or efficacy within the skin^27^. However, our data supports a need for broader pharmacokinetics and pharmacodynamics studies which include sites of less immediate clinical significance, including asymptomatic skin, if there is an intention to minimise the likelihood of patient relapse and/or the generation of drug resistance in parasites^27^.

Our third key finding is the extreme malleability of *Leishmania* dissemination pathways in response to immune perturbation, with re-infection of immune mice promoting dissemination from the skin to visceral tissues. Although we have not formally addressed skin-specific protection in this model, the reduction in visceral parasite burden because of prior infection suggests protective immunity has been established. Whether skin-resident CD4^+^ T cells play a role in skin immunity in this model, as has been shown for *L. major*^70,71^, remains to be determined. Nevertheless, this counter-intuitive re-routing suggests that the local immune response to secondary infection triggers the export of parasites from the skin back to the viscera. Hence, the skin has the capacity to replenish parasite diversity in visceral tissues and this indicates that dissemination patterns are not hardwired. Whether other non-cognate inflammatory responses act similarly to re-challenge is yet to be determined, but we speculate that the widespread heterogeneity in dissemination highways observed between individual mice could be a biological manifestation of an individual host’s immune history (i.e. the unique accumulation of inflammatory events, microbial dysbiosis, or co-infections^72^). Divergent dissemination patterns may offer an explanation for patient-to-patient variation in treatment response and propensity for relapse and imply that successful targeting of persistent parasite populations to avoid these negative outcomes could require an individualised treatment approach.

Our study has some limitations. Experiments were conducted using only female mice to avoid the potential confounding effects of inflammation caused by male aggressiveness. Hence, we cannot comment on whether dissemination patterns may exhibit sex-determined differences. Further, our experimental design analysed only a limited area of the skin for each mouse, and this may underestimate the full breadth of dissemination to, from and within the skin. Moreover, the invasive nature of current tissue sampling techniques means longitudinal tracking to determine whether “super-seeder” status is transient or a permanent feature of specific sites is currently not possible. Our analytical pipeline utilises an inference-based metric and cannot definitively identify migration events. We have thus combined this probabilistic inference of direction with network MST analysis and other ecologically-informed approaches to uncover key elements of dissemination in vivo.

In conclusion, our network-based analysis has provided an ecological framework for the investigation of parasite dissemination in experimental VL. Further, this approach has applicability more broadly for the study of other infectious (and potentially non-infectious) diseases. By revealing the skin as a dynamic reservoir capable of re-seeding visceral tissues and showing how this network is re-routed by immune perturbation, we provide new insights that can be integrated into strategies for preventing relapse and controlling the spread of *Leishmania* and emerging drug resistance.

## Methods

### Ethical approval

All animal experiments received approval from the University of York Animal Welfare and Ethical Review Body (AWERB), were run in accordance with the Animals (Scientific Procedures) Act, 1986 and were conducted under Home Office Project Licence (#PP0326977; “Immunobiology of leishmaniasis”).

### Mice

All studies used female C57BL/6J mice bred in-house at University of York, aged between 9 and 12 weeks old (barcode infections) or 15-17 weeks old (tdTomato infections). Parasites (see below) for mouse infections were freshly obtained from the spleens of long-term infected B6.*Rag2^−/−^*CD45.1 (RAG) mice as described previously^17^. For infections using the barcoded library, 3×10^7^ *L. donovani* amastigotes were injected intravenously in 100μl RPMI. For “re-infection”, mice were infected with 4-5×10^7^ stationary phase promastigote luciferase-expressing *L. donovani*. For confocal imaging experiments, mice were infected with 8×10^7^ amastigotes of the tdTomato-expressing *L. donovani* strain.

### Ex vivo luciferase imaging

Tissues were imaged ex vivo immediately after removal from mice post-cull. Skin was shaved post-mortem and Veet hair removal cream was then applied for 10 minutes, before washing with water to remove residual hair. Skin was scraped prior to imaging to remove adipose tissue.

For ex vivo imaging, 30mg/ml Firefly D-luciferin (Syd Labs: MB000102-R70170) in 1X PBS was applied to the surface of tissues, then incubated for 5 minutes in the dark. For skin, luciferin was applied to the hypodermal side. Imaging was performed with an IVIS Spectrum bioluminescence imaging system (PerkinElmer) with an open filter. Light detection was captured via a charge-coupled device (CCD) camera. Exposure was for up to 5 minutes for imaging and images were obtained and processed using Living Image version 4.7.4.21053; this constituted drawing regions of interest around each tissue to quantify flux for each sample. This data was then further analysed in R, by quantifying the background rate of flux for each tissue type using the mean flux obtained from naïve tissues; this background was subtracted from the flux for each infected tissue sample, to give a total flux above background level value for each tissue sample. Any negative values were set as 0. These values were then changed to a log scale, with any undefined values changed to 0.0000001, for the purposes of visualisation and statistical analysis.

### Confocal imaging in skin

For confocal imaging of mouse skin, the skin was shaved post-mortem and Veet was applied for 10 minutes, before washing with water. Mice were skinned, adipose tissue was removed and skin was imaged from the hypodermal side using an LS980 Upright Microscope (2.5X magnification; pin hole set to 500μm) using Zen version 3.9. Spectral fingerprinting based on a purified amastigote tdTomato *L. donovani* sample was used to generate a spectral profile for tdTomato, whereas an autofluorescence signal was derived from uninfected age matched control skin (**Supplementary Figure 3**). Image processing was performed in Zen Blue Version 3.9.101.02000.

### In vitro infection of bone marrow derived macrophages

BMDMs were incubated with 5X multiplicity of infection (MOI) promastigote parasites at 37 LJC for 1 or 3 hours. Slides were fixed with methanol for 10 minutes and stained with 10% Giemsa stain for 3 minutes. The parasite burden per cell was then assessed by counting by eye for 100 host cells per condition. Poisson distributions were fitted to the parasite count per cell data for each barcoded parasite infection experiment at each time-point using the “fitdist” function in the fitdistrplus package in R^73^; scripts are available on Github.

### Parasites

All *L. donovani* lines used were generated from the Ethiopian strain (MHOM/ET/1967/HU3), also referred to as LV9 or HU3. The *L. donovani* T7 Cas9 line used for the generation of the barcoded library was generated following the method described in^74^ for *L. infantum*. The luciferase-expressing parasite line *L. donovani* RE9H LV9 encoding a red-shifted firefly luciferase gene (Ppy-RE9H) in the ribosomal RNA locus^75,76^ and a tdTomato-expressing *L. donovani* strain^77–79^ were generated as described. Promastigote parasites were cultured at 25 LJC in SDM-79 media (Gibco: 074-90916N), supplemented with either 10% or 20% (for transfection recovery only) heat-inactivated FCS (Gibco: A5256701), with 5 μg/ml hemin (Sigma-Aldrich: 51280-5G), 10 μM 6-biopterin (Sigma-Aldrich: B2517-25MG), 100 U/ml penicillin and 100 μg/ml streptomycin.

### Development of the barcoded parasite library

Genome editing to insert the unique 20bp barcodes was performed in the *L. donovani* strain constitutively expressing Cas9 and T7 RNA polymerase, generated following the protocol described in^74^. The 20bp barcode sequences were incorporated into the exogenous HYG locus used for insertion of the T7 and Cas9 expression cassette using reverse primers. The forward primer used was consistent for all insertions (OL12845). Two single guide RNAs (sgRNA) targeting the 5’ and 3’ insertion region within HYG were designed using the Eukaryotic Pathogen CRISPR guide RNA/DNA Design Tool (http://grna.ctegd.uga.edu). sgRNA templates and the repair cassettes (carrying a blasticidin resistance marker) were generated by PCR, using the CRISPR-Cas9 toolkit for kinetoplastids^80–82^. All primer sequences are provided in **Supplementary Table 17.**

PCR products were purified using the QIAquick PCR Purification Kit (Qiagen: 28104). For transfection, sgRNA and the repair cassette was co-transfected into 2×10^6^ promastigotes using the P3 Primary Cell 4D-Nucleofector X Kit L (Lonza: V4XP-3024) with program FI-115. After electroporation, cells were immediately transferred to pre-warmed SDM-79 media, containing 20% FCS. Parasites were incubated at 25 LJC with 150 μg/ml blasticidin initially, which was then reduced to 75 μg/ml for first passage and to 20 μg/ml for maintenance culture. Genomic DNA from recovered parasites was extracted for diagnostic PCR and successful barcode integration was visualised using a product size-based PCR (approximately 2 Kbp), using OL12882 and OL12884 primers (see **Figure 1a**). Sanger sequencing of the initial 10 barcode insertions was also performed for validation.

### Barseq sequencing and analysis

Tissue samples were flash frozen on dry ice and stored at −80 LJC. For bone marrow, the bones were flushed using un-supplemented RPMI with a 26G needle; marrow was then centrifuged at 3100 rpm for 10 minutes at room temperature. The DNA was extracted using Qiagen DNeasy Blood and Tissue Kits (Qiagen: 69506), following the tissue DNA extraction instructions for all samples except bone marrow and isolated or cultured parasite samples. For these two sample types, the cell culture instructions were followed. The 56 LJC digestion step was performed for 10 minutes for the bone marrow and parasite samples, for 1-2 hours for all tissue samples except skin, and overnight for skin.

For sequencing, the barcodes were amplified using a two-step PCR approach, before addition of Nextera XT Indexing primers. For the first PCR step, the amplification PCR used OL12833 and OL13833 and 1 unit Verifi Mix (PCR Biosystems: PB10.43-05), 0.4 nM 10 nm gold nanoparticles in citrate suspension (Sigma-Aldrich: 741957) and 0.025 mM MgCl_2_ (Sigma-Aldrich: M1028) in a 25 μl reaction, for 30 amplification cycles, with an annealing temperature of 60 °C and an extension time of 40 seconds. The first PCR product was then cleaned using 0.9X Agencourt AMPure XP beads (Beckman Coulter: A63882; see below). For the second PCR step, amplification of the purified product used OL14195 and OL14196 and 1 unit Verifi Mix (PCR Biosystems: PB10.43-05), 0.4 nM 10 nm gold nanoparticles in citrate suspension (Sigma-Aldrich: 741957) and 0.025 mM MgCl_2_ (Sigma-Aldrich: M1028) in a 25 μl reaction, for 6 amplification cycles, with an annealing temperature of 65 °C and an extension time of 40 seconds.

Samples were cleaned again using 0.9X Agencourt AMPure XP beads (Beckman Coulter: A63882). For the purification (both steps) samples were incubated with the beads at room temperature for 5 minutes. They were placed on a magnetic rack to clear, followed by removal of supernatant. Beads were then washed twice with 75% ethanol and dried for approximately 3 minutes before elution, using Molecular Biology Grade Water (ThermoFisher: AM9932) for 5 minutes at room temperature. For the library preparation, PCR products underwent 9 PCR amplification cycles with Q5 Polymerase 2X Master Mix (New England Biolabs: M0492L), following the manufacturer instructions. Sample purification was performed using 0.9X Agencourt AMPure XP beads (Beckman Coulter: A63882), eluted into a low TE buffer, and pooled samples were then sent to Azenta Life Sciences for 150 base paired end sequencing using an Illumina Sequencer.

### MATLAB simulation

To assess the accuracy of STAMP FP estimation for different sizes of barcode library (**Figure 1b** and **1c**), we used MATLAB to simulate barcoded “parasites” passing through a range of bottlenecks. An input pool of barcoded “parasites” was generated as a matrix of 5×10^7^ individuals, with an even number allocated to each barcode, for barcode totals ranging from 10 to 500 in total. Bottlenecks were simulated as a probability of an individual parasite “passing” the obstacle, ranging from 0.0000001 to 0.5. The simulation was repeated for 100 “experiments” for each bottleneck and barcode total combination.

We initially ran a basic simulation assuming all barcoded parasites replicated at the same rate post-bottleneck, by counting the individuals with each barcode after the bottleneck was applied. We then refined the simulation to reflect a 5% random variation in overall growth rates, by multiplying the total count of individuals post-bottleneck for each barcode by a randomly generated number between 0.95 and 1.05.

For the simulation of clonal expansion, the same procedure as the 5% random variance simulation was followed, but immediately after the bottleneck, we randomly selected 1 in 100 (common), 1 in 10000 (medium) or 1 in 1000000 (rare) individuals to ‘clonally expand’, by multiplying their count by 10 to simulate a growth rate for these individual parasites of 10x the average. We then used the counted total of all passing ‘parasites’ as our ‘true FP’ and computed FP for our ‘STAMP-estimated FP’, using the method defined by Abel et al^43^, based on original equations from Krimbas and Tsakas^45^. The equation used was as follows:

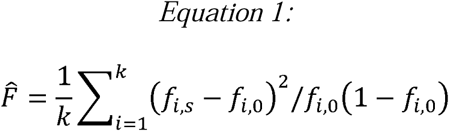

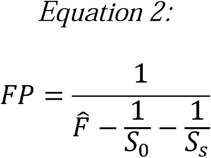

where *f_i_*is the frequency of barcode *i,* at input (*f_i, 0_*) and after the bottleneck (*f_i, S_*), *S_0_*and *S_S_* are the total parasite counts for each, and *k* is the total number of barcodes. The original equations contain a generation term, *g*, which for our analysis we set as 1. The accuracy of the estimation was then assessed via Kendall correlation, using the cor.test function in R. The MATLAB script for the simulation and R script for the analysis are available in Github (see Data and Code Availability section for details).

### Barcode quantification

Barcode sequence counts were extracted using a Rust script (available on Github; see Data and Code Availability section for details). This script used the primer sequence directly before the barcodes to pull out 20bp barcode sequences from all forward reads. Barcodes with each sequence were counted and those with 500 or fewer reads were then merged with the barcode above the 500 read threshold from which they each had the lowest Levenshtein distance. If they were not within 5 Levenshtein edit distances of any barcodes above the 500 read threshold, they would not be merged for counting. An R script (available on Github) which pulled out only the canonical barcode sequences from this list of counts was used to generate the analysis matrices of barcode counts for all samples analysed. Prior to subsequent analysis, the barcode with sequence TGGATCTTAGGACGCAACAT was removed due to indications in preliminary studies (Loughrey, unpublished^83^) that it exhibited an usually high abundance in an unexpectedly large number of tissue samples across multiple experiments.

### Diversity analysis

Within tissue parasite diversity was quantified using the Shannon Index, calculated using the Vegan package in R^84^. The Shannon Index was quantified as:

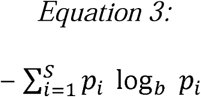

where *p_i_* is proportion of “species” *i* (in this case barcodes) and *S* is total number of “species” (barcode total) and *b* is the default logarithm base option in Vegan (natural logarithm *e*), as defined by Hill^60^. Analysis scripts are available on Github (see Data and Code Availability section for details).

### Genetic Distance analysis

GD (*D_ch_*) was calculated as Cavalli-Sforza chord distance^44^ as shown:

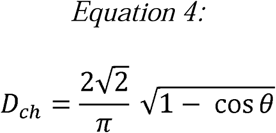

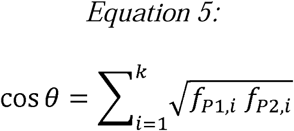

where *f_i_* is the frequency of barcode *i* in tissue parasite populations *P1* and *P2* and *k* is total number of barcodes. When using GD to build full networks, we used GD to initially structure the networks, then used 1-GD (i.e. genetic relatedness) for edge weights for visualisation. All GDs of above 0.5 were removed. This calculation and network generation was done using custom R code available on Github (see Data and Code Availability section for details).

### Founder Population metrics and network analysis

The evidence we had of serial bottlenecks and a highly interconnected network of parasite populations led us to speculate about the direction of parasite movements between tissues, and as such we decided to calculate FP from tissue sources to tissue recipients for all pairwise comparisons within each individual mouse (see **Figure 4a**). We termed this directional metric Tissue-sourced FP (TSFP). Before calculating this value, we removed all samples with total barcode reads of below 50000, as samples below this cut off confounded the calculation due to the large difference between *S_0_* and *S_S_*; removed samples are noted in **Supplementary Table 18**. Sample barcode counts were then filtered for each individual comparison pair in order to remove all barcodes with a count of 0 in the source tissue from both samples being analysed. TSFP was calculated based on the STAMP method^43,45^ as described below:

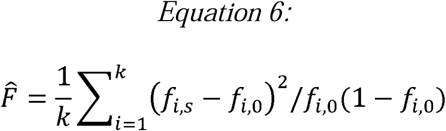

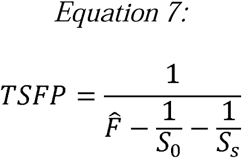

where *f_i_*is the frequency of barcode *i,* in the input source tissue (*f_i, 0_*) and in the tissue recipient (*f_i, S_*), *S_0_* and *S_S_* are the total barcode reads for each sample, and *k* is the total number of barcodes included in the analysis.

To calculate the overall directional movement of parasites, we found the difference between TSFP in each direction for each tissue pair and used the positive value to build a network of directional parasite movements. We termed this value Fractional FP (FFP). For network construction and analysis, we used the igraph package in R^85,86^, with ggraph^87^ used for generating the final figures. FFP was used as the edge weights for building full networks. For computation of minimum spanning trees (MSTs), we used 1/FFP as edge weights. In both cases, we removed all edges with FFP below 20. For analysis of degree (in and out), we used the igraph ‘degree’ function on the full network. For eigen centrality analysis, we used the ‘eigen_centrality’ function in igraph on the undirected full graph. All scripts are available on Github (see Data and Code Availability section for details).

### Validation

To validate our analysis of TSFP and FFP, we first assessed the likelihood of false migration detection in randomised datasets. To do this, we took our original barcode count matrix and randomly reallocated the counts for each barcode to another barcode in our list, across all samples in the dataset. We repeated this for three simulated datasets in total (**Supplementary Tables 19-21**). We reasoned that this approach preserved the number of barcodes and count distributions for tissue populations of parasites, but randomisation meant they were not related to each other. We calculated pairwise TSFP and FFP for each “sample” and calculated the percentage of values which were above a set of threshold FFPs, ranging from 1 to 500 (**Supplementary Table 22**), using the AboveThreshold function in the Seurat R package^88–92^. We noted that the percentages above thresholds were very low in all three simulations when compared to our actual dataset of within-mouse FFPs.

We then used our INPUT samples to create a threshold value for confidence of FFP. We reasoned here that our populations within tissues cannot truly be sources of the INPUT pool, whilst the INPUT pool must be the ultimate source for at least part of the population within our tissue samples, although this could constitute a very small proportion of the end population under circumstances of very restrictive bottlenecks, high inter-tissue migration and/or selective pressures affecting replication dynamics. We therefore calculated TSFP and FFP in both a “true” migration direction, defined as INPUT pool (population A) to sample (population B) and a “back” migration direction, defined as sample (population B) to INPUT pool (population A) for all combinations of our samples and INPUT pools. **Supplementary Figures 4a and 4b** show the histograms for TSFP and FFP respectively. For visualisation on a log scale, we changed all FFP values below 1 to 0.9 and all negative TSFP values and any NA values to 0. Using these histograms, we defined our FFP threshold for our network and MST creation as those above 20 (log3 approximately), based on the shift between true and back migration distributions we observed. We additionally used this analysis to compare the true vs back migration above threshold for each tissue type (**Supplementary Tables 23** and **24**), as we hypothesised that this would vary by tissue due to underlying differences in both population structures and in relatedness to the original INPUT pool itself, due to initial colonisation dynamics. We confirmed this to be the case, meaning we cannot establish definitive “false positive” rates for our metric, as this will be affected by variables such as sample type and infection model used. Importantly we show here that whilst tissues such as spleen show relatively high “false” migration, as predicted, they show consistently higher levels of “true” migration. Of note in terms of our work’s conclusions, we also show that the skin shows no evidence of any “false” migrations above our threshold of 20. To further provide confidence in our analysis approach, we tested it on three published datasets^47,52,55^**(Supplementary Figure 5)**. From Hotinger et al^52^ (**Supplementary Figure 5a**), we used a dataset of multiple samples from 73 individual mice, with a set of individual inoculums for different infection groups, some of which had up to three replicate samples; in this case, we matched inoculum to the correct mice for our analysis. From Lebrun-Corbin^47^ (**Supplementary Figure 5b**), we used a set of multiple samples from 12 mice, with 30 replicate INPUT inoculum samples. From Hullahalli and Waldor^55^ (**Supplementary Figure 5c**), we used a set of multiple samples from 32 mice plus 5 INPUT pools as replicates of a single inoculum.

Whilst there are biological insights to these analyses, these are beyond the scope of this work. **Supplementary Figure 5** shows the thresholding histograms for these datasets, which we find illustrative of three key points. Firstly, that our analytical pipeline can be effectively implemented on a range of microbial population datasets, across diverse tissues, pathogens and infection model conditions. Secondly, that a skewed barcode distribution (such as that induced by the amastigote passage process our parasites underwent) does not seem to hinder, and in fact may be beneficial for, threshold determination, as evidenced by the clear distribution shift in our data (**Supplementary Figure 4**). Library size (which differed widely between the datasets) also does not appear to affect thresholding clarity; we hypothesise that the dynamics of within-host replication patterns may play a role in the ability to clearly discern directionality, given that Hullahalli and Waldor specifically report on clonal expansion occurrences in their dataset hampering FP calculations^55^. Thirdly, thresholds appear to vary between experiments. This is likely related to the biological aspects of the systems in question, given they all utilise different pathogen strains with divergent initial colonisation patterns and specific sample types measured. However, a smaller library size does not seem to reduce threshold discernment in itself.

We were cognizant that background similarity of populations is inherent in systems derived from a common ancestor population, and that we cannot directly measure the initial similarity of tissue populations upon initial colonisation within a single animal immediately after injection to account for this. This is an inherent limitation of population dynamics for the study of dissemination in vivo, regardless of metric choice. The only scenarios in which it can be completely avoided are one in which there is no intermixing or migration to measure to begin with combined with a very high level of initial diversity or very restrictive initial bottlenecks to individual tissue colonisation (i.e. no overlaps in barcode presence between tissues at all), in which case there is no dissemination to measure, or alternatively by using a system in which barcode expression or editing is induced in situ after initial colonisation.

Therefore, we also used our dataset to calculate TSFP and FFP for a matched number of randomly selected comparisons of samples from disparate mice and compared these to our within-mouse values for each metric (**Supplementary Figure 6**). When comparing density distributions for all values above our threshold of 20, we observed that, as expected, background similarity of populations generates an inference of migration. However, we note a sustained shift in the distribution of the within-mouse comparisons towards increased TSFP (**Supplementary Figure 6a**) and FFP (**Supplementary Figure 6b**), giving us confidence that we were detecting true dissemination dynamics. This relatedness confounder likely differs between model systems depending on initial bottleneck events, population structural differences inherent to different niches and with the relative complexity of inter-seeding patterns. Importantly, it will apply even in experiments with larger library sizes (although it may be missed in these cases, if using between animal comparisons for detection) and with the use of pre-established population dynamics metrics such as Cavalli-Sforza chord distance. Therefore, experimental design should incorporate time-based comparisons between groups, interpretations should be informed by biological knowledge of the system in question and of biologically plausible relationships, whilst analysis approaches should consider using a combination of metrics with network analysis, including MSTs, for more robust determination of dissemination pathways.

All datasets and R scripts for this validation are supplied on Github (see Data and Code Availability section for details).

### Experimental design and statistical analysis

Only female mice were used to avoid non-specific tissue damage in the skin caused by fighting in cages of male mice. One mouse was excluded from further analysis after dissection due to an absence of hepatosplenomegaly (which would be evidence of established infection at 4 weeks p.i). One mouse was killed early due to welfare concerns and was excluded from subsequent analysis.

Samples with fewer than 50000 reads for canonical barcode sequences were removed from TSFP and FFP analysis (but were used for SI and GD analysis). Sample read counts are supplied in **Supplementary Table 22**. Statistical tests for TSFP (including GLMM fitting) were performed on data after removal of samples with fewer than 50000 read counts. Animal experiments were performed using the same pool of barcoded amastigotes extracted from passage mice. Differences in the barcode frequencies between pools derived from separate mice that arises from “bottlenecks” associated with passage ruled out amalgamation of replicate experiments and direct comparison for Barseq-based statistical analysis. Sample sizes are given in Results and/or Figure legends. The “single infection” group used for Barseq statistical comparison to “re-infection” comprised 5 mice which were infected for 12 weeks and were also used in the “chronic” infection group; the remaining three mice in this group were not used for comparison to “re-infection” as they were infected for 10 weeks. For Barseq analysis for each mouse, a single sample was sequenced for liver, spleen, gut and lung and for two separate inguinal lymph nodes and femur/tibia. For skin 6-12 biopsies were taken per mouse as indicated in Results and/or Figure legends.

Data processing, statistical analysis and figure generation were performed using the tidyverse^93^, ggraph, ggpubr^94^, igraph, fitdisrplus, naniar^95^, Seurat and rstatix^96^ packages in R version 4.4.3. Mann-Whitney-Wilcoxon tests (two sided) were used for in vivo parasite population comparisons of diversity (Shannon Index), Genetic Distance and TSFP where indicated in figure legends.

We used a Compound Poisson GLMM to analyse the fixed effect of time p.i (early vs late) on parasite diversity (Shannon Index) across all skin biopsies for each mouse, with mouse ID as a random effect. All GLMMs were fitted and compared using the cplm and tweedie packages^97–100^ in R. We compared models with both effects and with only mouse ID or only time p.i using the “gini” function in cplm. This approach fits a Lorenz curve and calculates the Gini index for each model fit against this curve; selecting the model with the lowest maximal absolute Gini index when compared against all other models thus gives us the closest model fit to our dataset^101^.

We likewise used a Compound Poisson GLMM to compare models for Genetic Distance between visceral tissues and skin biopsies, and for model comparisons for TSFP to viscera from skin biopsies and vice versa. For these models, we used time p.i (Genetic Distance only) and reinfection vs single infection (TSFP only) plus visceral tissue identity as the fixed effects and mouse ID again as the random effect. Model comparison was performed in the same manner as described above, but models tested included the interaction of the two fixed effects plus the random effect, the same model but without interaction of fixed effects, and then a range of models with each effect removed. All GLMM model outputs and comparisons are included in the **Supplementary Note**.

## Data and code availability

Summary statistical data and raw read counts for barcode abundance are available both as supplementary data files with this manuscript and in the Github repository at: https://github.com/CiaraLoughrey-scientist/barseq-Leishmania-2025. Other analysis scripts are also provided in the Github repository. Raw Illumina sequencing data is available in NCBI SRA database with Project ID PRJNA1392034 (http://www.ncbi.nlm.nih.gov/bioproject/1392034). Raw image files for tissue and skin imaging are available on request. Schematics were created using Biorender.

## Acknowledgements

The authors thank Natalie Prow and Najmeeyah Brown for assistance with animal experiments, Ewan Parry for advice on primer design for Bar-seq, Charlotte McNiven for help with parasite culture, João Luís Reis Cunha for advice on code queries and figure creation, the University of York BSF for their animal husbandry and Katherine Newling for designing the barcode sequences for the library.

The work was funded by Wellcome Investigator Awards to PMK and JCM (#224290 and #223045 respectively; https://wellcome.org). CL was supported by a Hull York Medical School PhD studentship.

## Author Contributions

C.L., P.M.K. and J.C.M conceived of and designed the study. P.M.K. and J.C.M. obtained funding. C.L., J.C.M. and P.M.K wrote the manuscript and all authors reviewed and provided feedback on the manuscript. C.L. and J.P. designed the simulation. C.L. and J.B.T.C. created the barcoded parasite library. C.L. performed the in vitro experiments. C.L., H.A. and Sh.De. performed animal experiments. C.L. performed tissue processing. C.L., S.J., L.G. and Sa.Do. optimised protocols for and generated the Barseq amplicon libraries. C.L., G.C. and K.H. created the confocal imaging protocol and performed the confocal imaging. C.L. analysed the data and created the figures, with oversight from P.M.K and J.C.M. J.P., Sh.De., and A.D. provided input and support in data processing, analysis and code development.

---

**Extended Data Figure 1:**
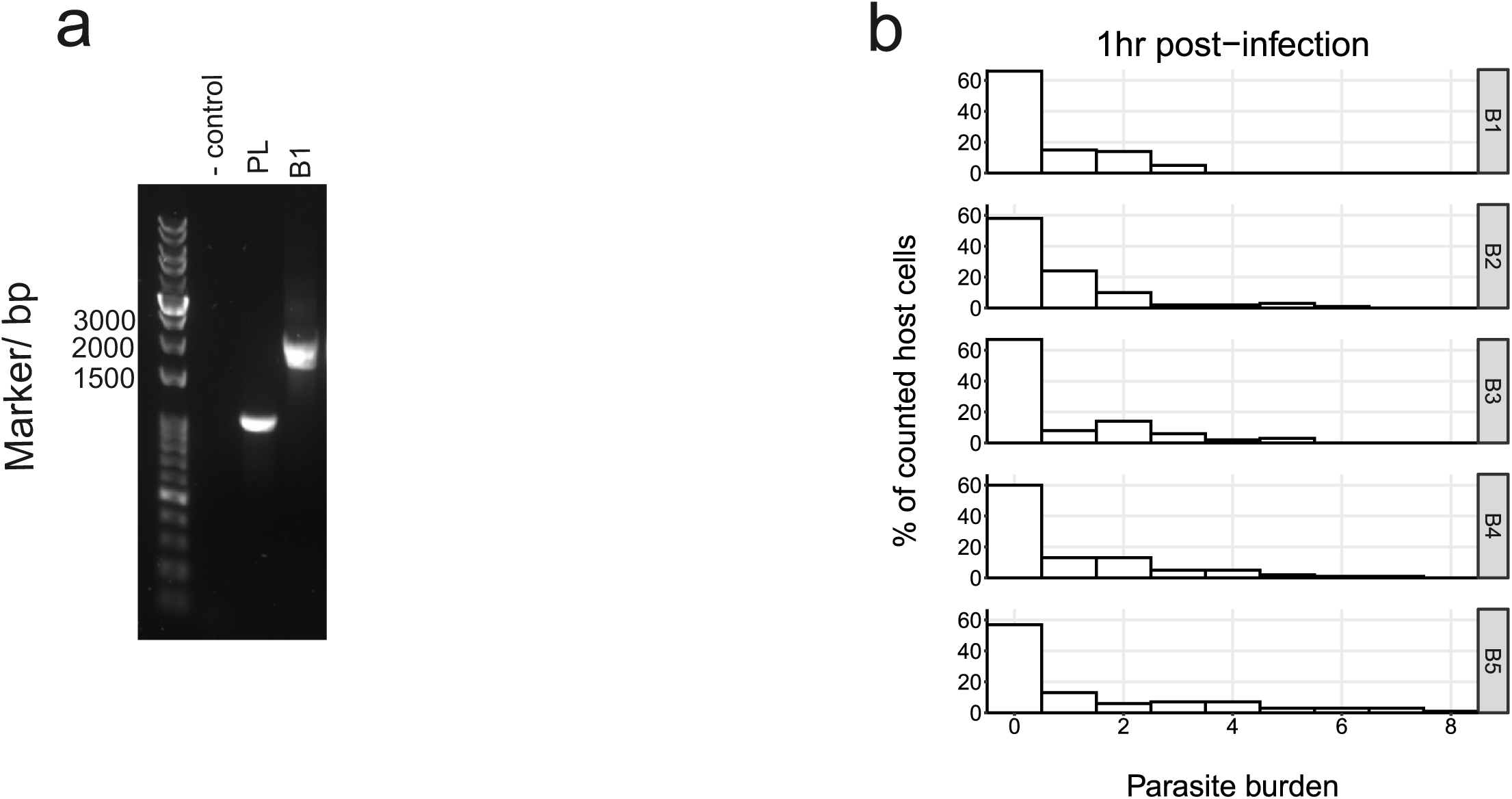
Barcoded parasite validation. **a,** Validation PCR for barcode insertion in *L. donovani*. PL = parental line with HYG present. B1 = mutant parasites containing barcode 1 (B1) and BSD selection marker. b, Histograms of in vitro parasite infection burdens after a 1 hr incubation of promastigote parasites with murine bone marrow derived macrophages for five individual barcoded parasite lines, here termed B1-5.

**Extended Data Figure 2:**
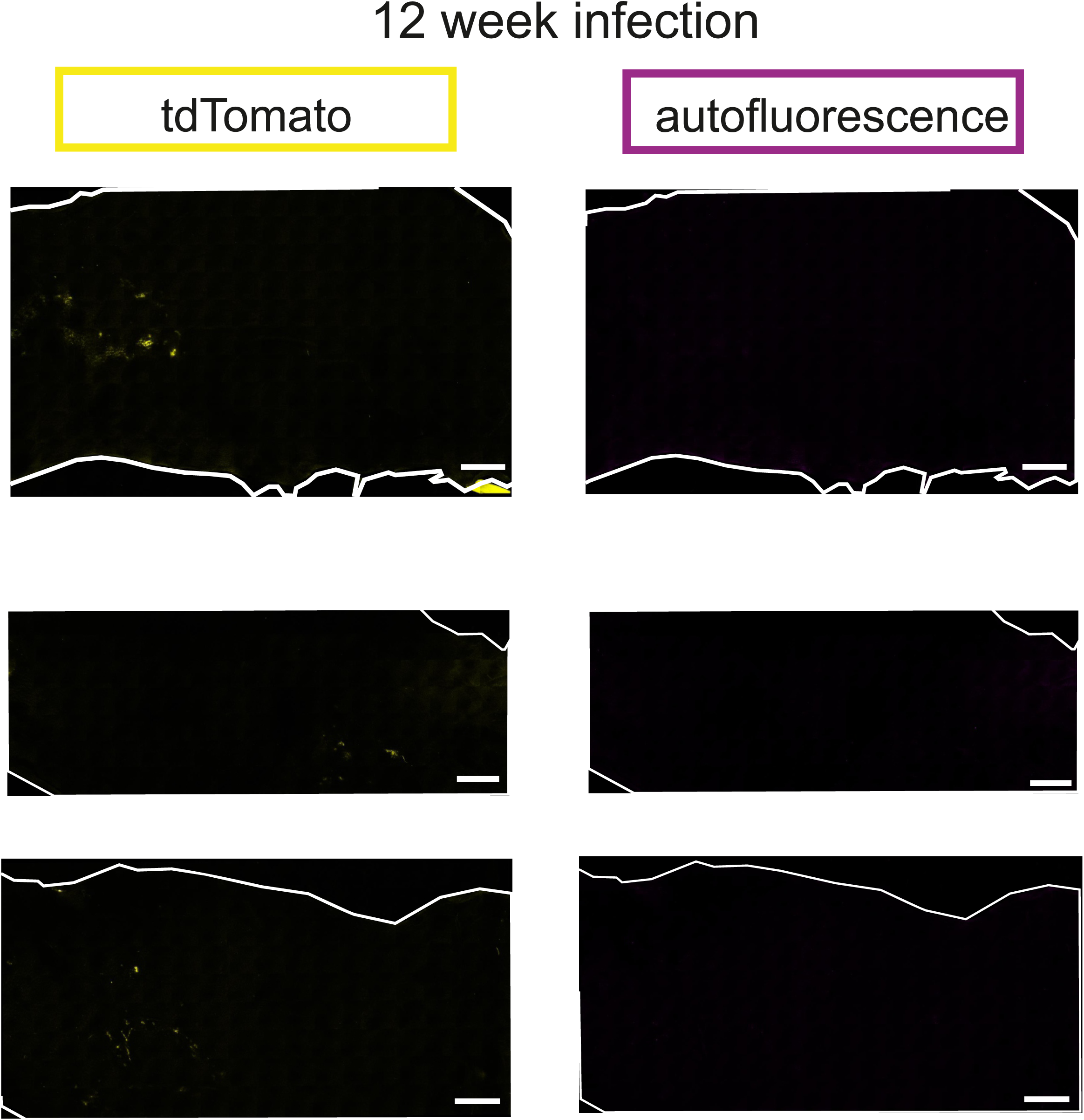
Detection of tdTomato-expressing *L. donovani* in the skin of C57BL/6J mice. Whole skin images (after stitching of panels) of infected skin at 12 weeks post-infection (n=3 female mice). tdTomato signal shown in yellow. Autofluorescence signal shown in magenta. Scale bar represents 5000μm.

**Extended Data Figure 3:**
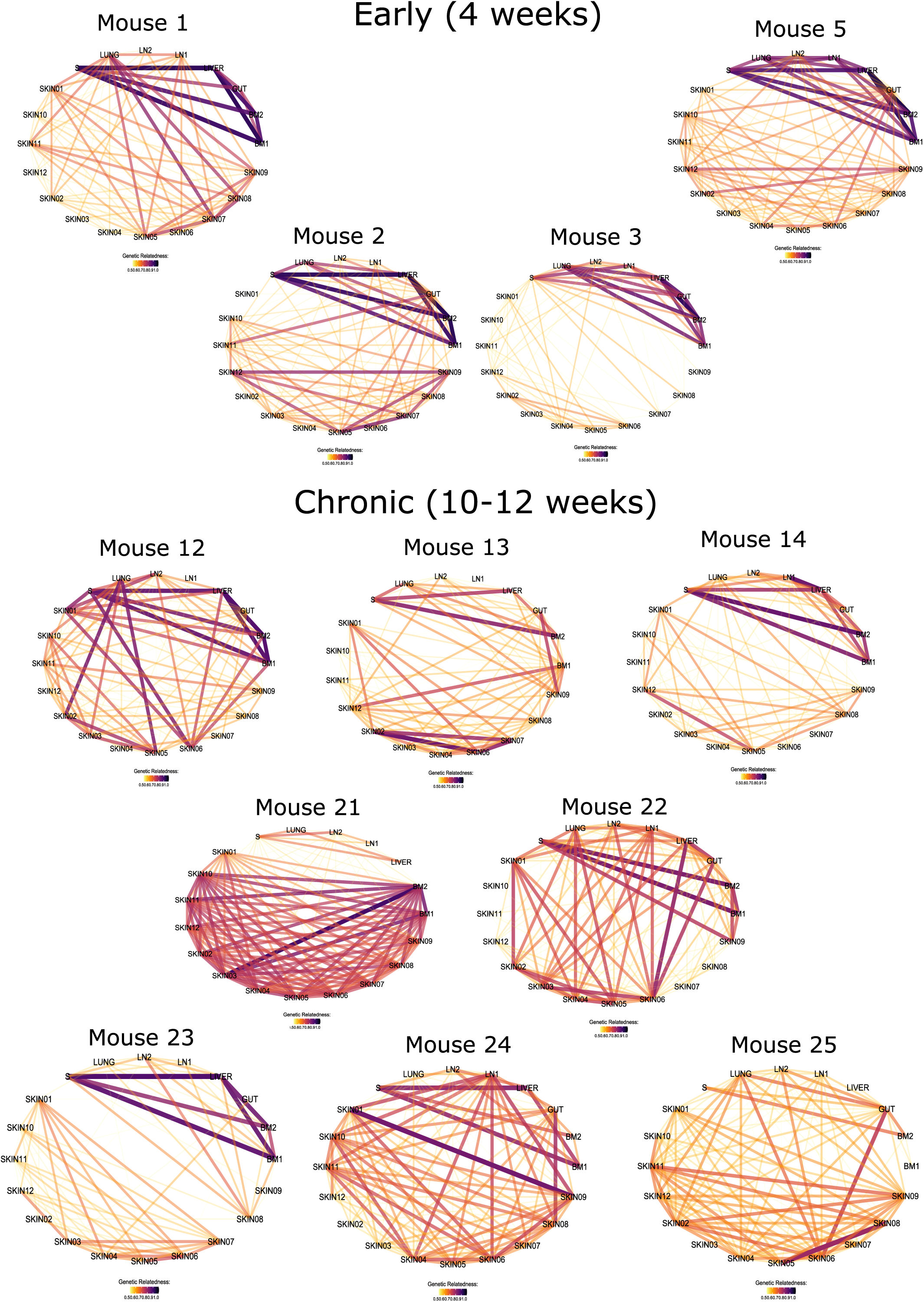
Tissue parasite population connectedness networks. Network representations of GD (edges) between parasite populations in each tissue (vertices). Networks for all mice at 4 weeks (early) and 10-12 weeks (chronic) post-infection, with all GD above 0.5 removed. Line colour indicates genetic relatedness (1-GD) as shown in the legends for each network. Thicker, more opaque connecting lines indicate a higher relatedness (lower GD).

**Extended Data Figure 4:**
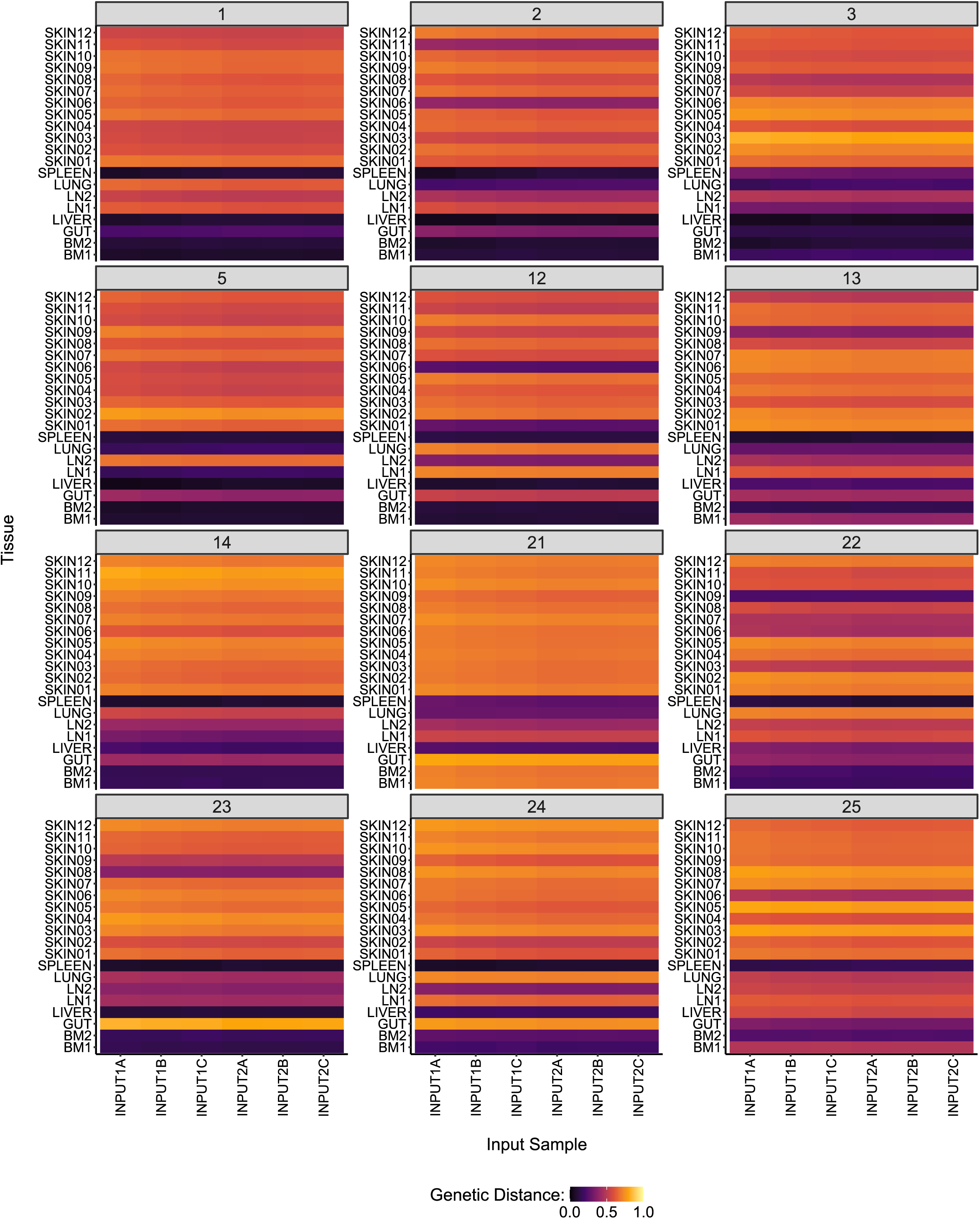
Tissue parasite population relatedness to input. GD of parasite populations to input pool (n=6 sequencing repeats from a single culture) in all tissues for all mice at 4 weeks (early) and 10-12 weeks (chronic) post-infection. Strip text identifies mouse ID. Colour scale applies to all heatmaps.

**Extended Data Figure 5:**
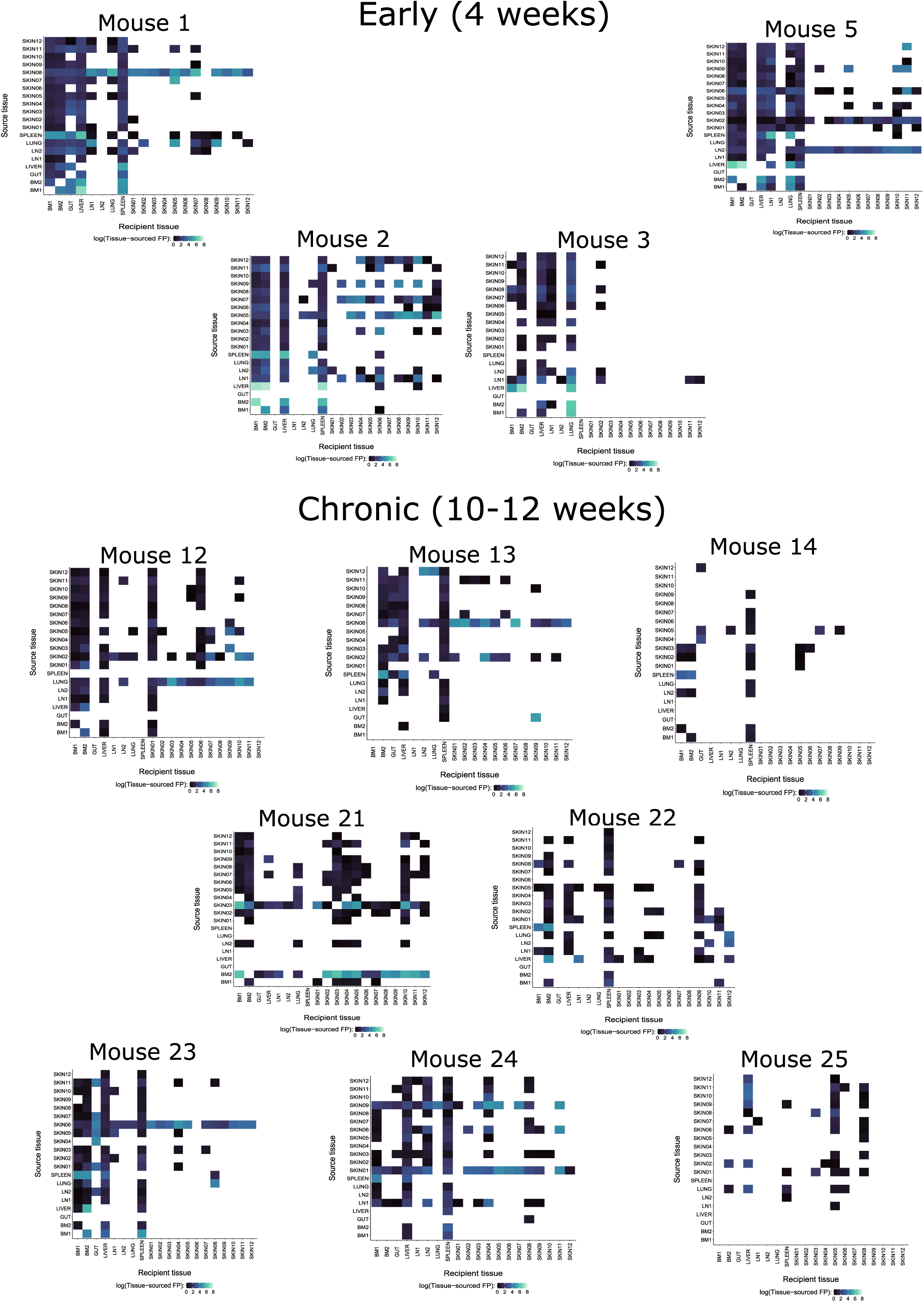
Tissue to tissue parasite seeding heatmaps. Scale represents log(TSFP) for each directional tissue to tissue relationship. White indicates those TSFP values < 1 or tissues which were excluded in filtering. No further thresholding was applied to TSFP for the heatmaps shown.

**Extended Data Figure 6:**
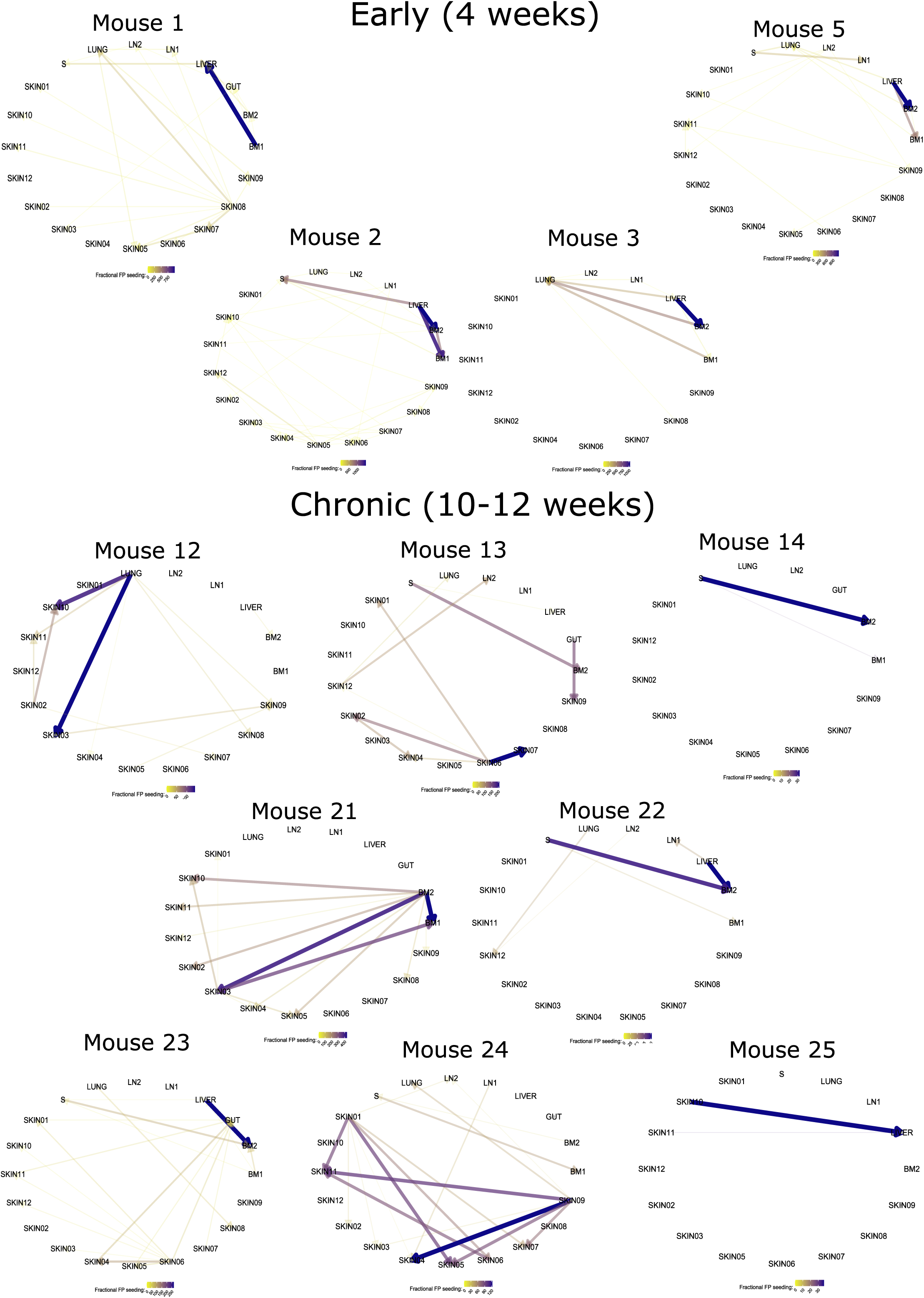
Tissue to tissue parasite seeding networks. Full networks constructed using FFP as the metric for edge weight for all mice at 4 weeks (early) and 10-12 weeks (chronic) post-infection. Line colour indicates FFP as in legends. Thicker, more opaque lines indicate a higher FFP in the direction indicated by the arrowhead. A threshold

**Extended Data Figure 7:**
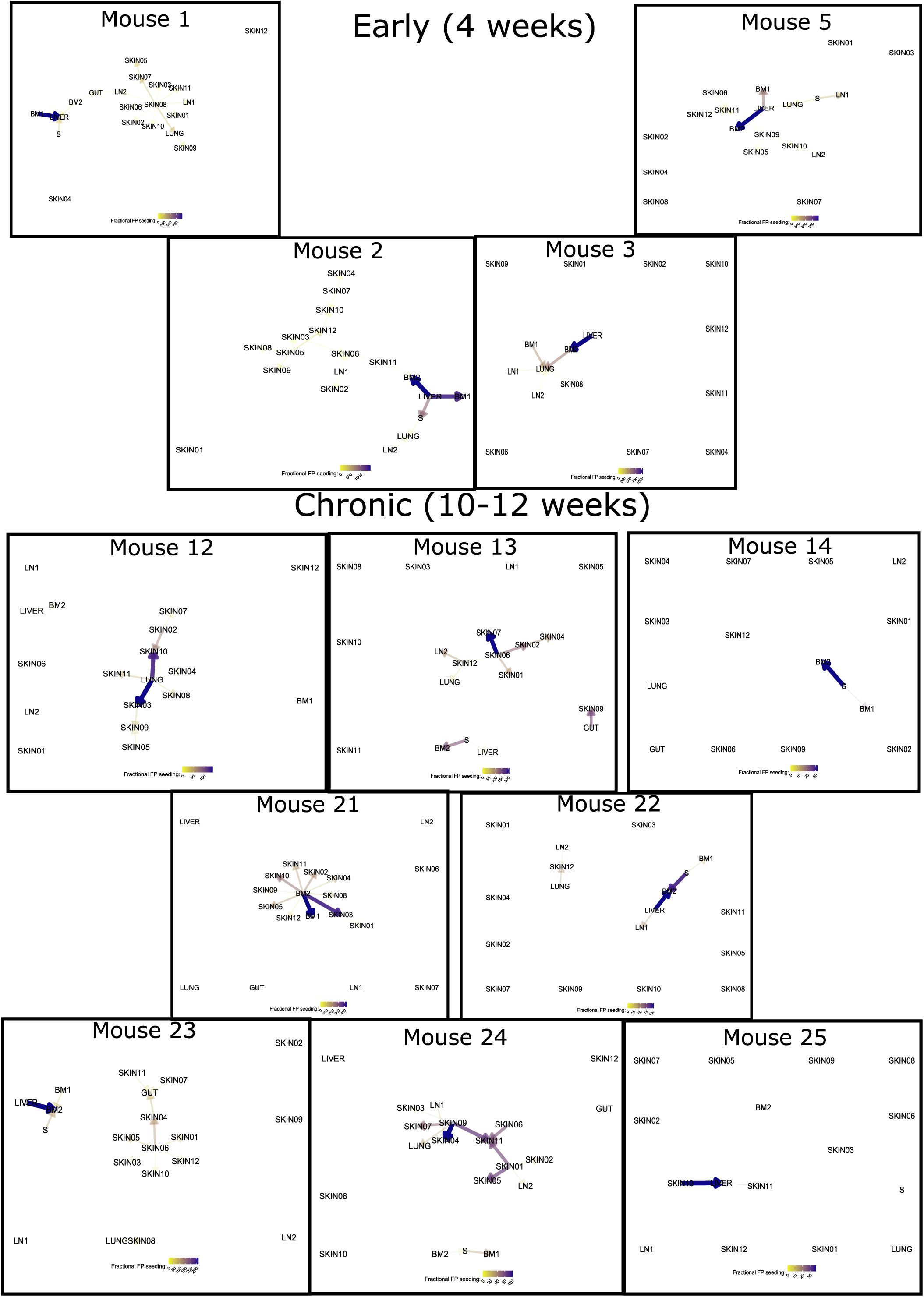
*In vivo* parasite dissemination “highways”. Minimum spanning tree (MST) network representations of FFP for all mice at 4 weeks (early) and 10-12 weeks (chronic) post-infection. Legends, line colour and thickness indicate FFP. Arrowheads represent direction of FFP. A threshold of FFP > 20 was applied for initial network creation, prior to MST computation.

**Extended Data Figure 8:**
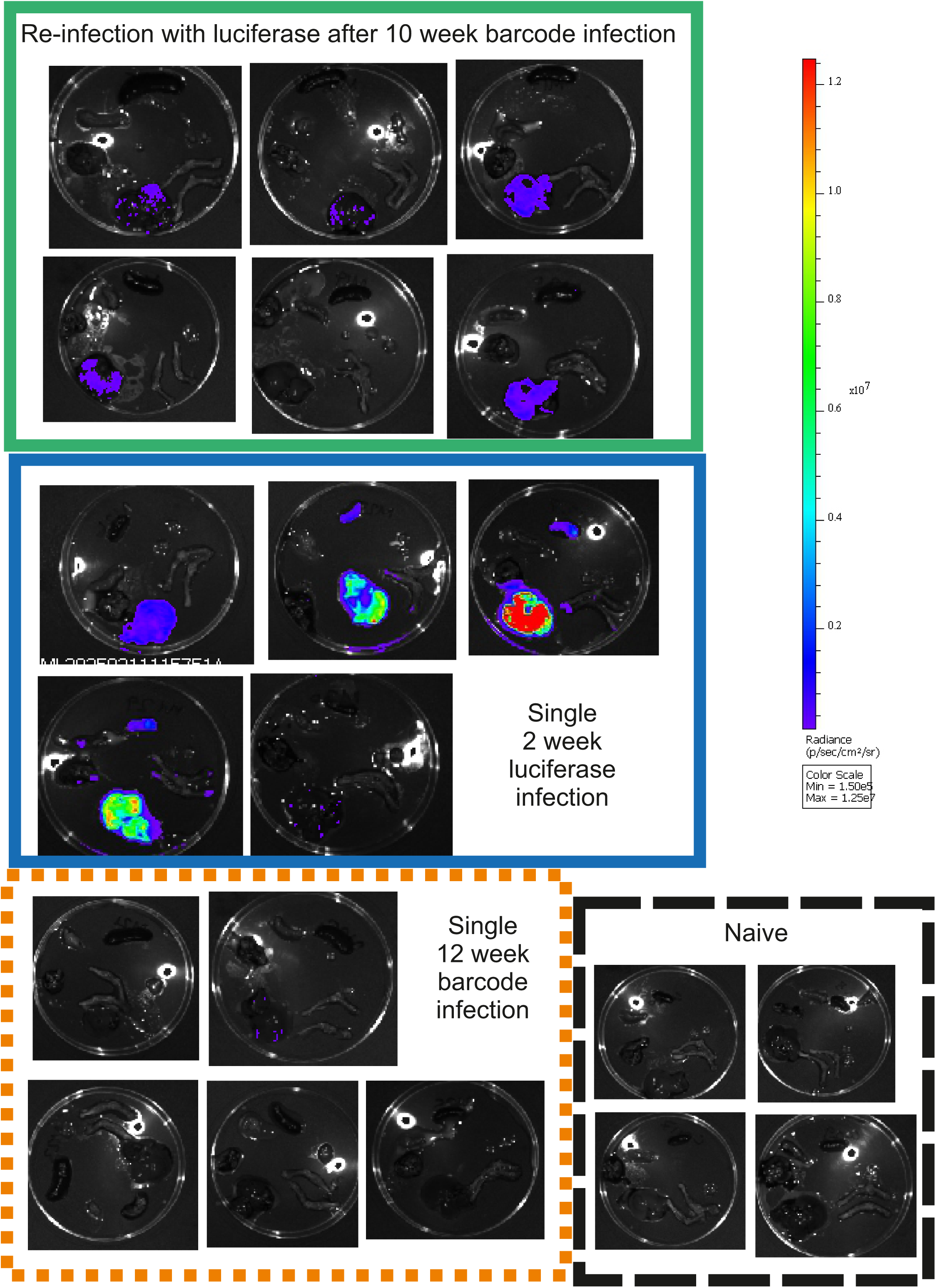
Tissue parasite burdens ex vivo. *Ex vivo* IVIS imaging of tissues from each individual mouse, with infection group as indicated by the box label. Radiance scale (p/sec/cm^2^/sr) applies to all images.

