## Supplementary Figure 1 for "Bi-directional highways and super-seeder tissues underpin parasite dissemination in experimental visceral leishmaniasis"

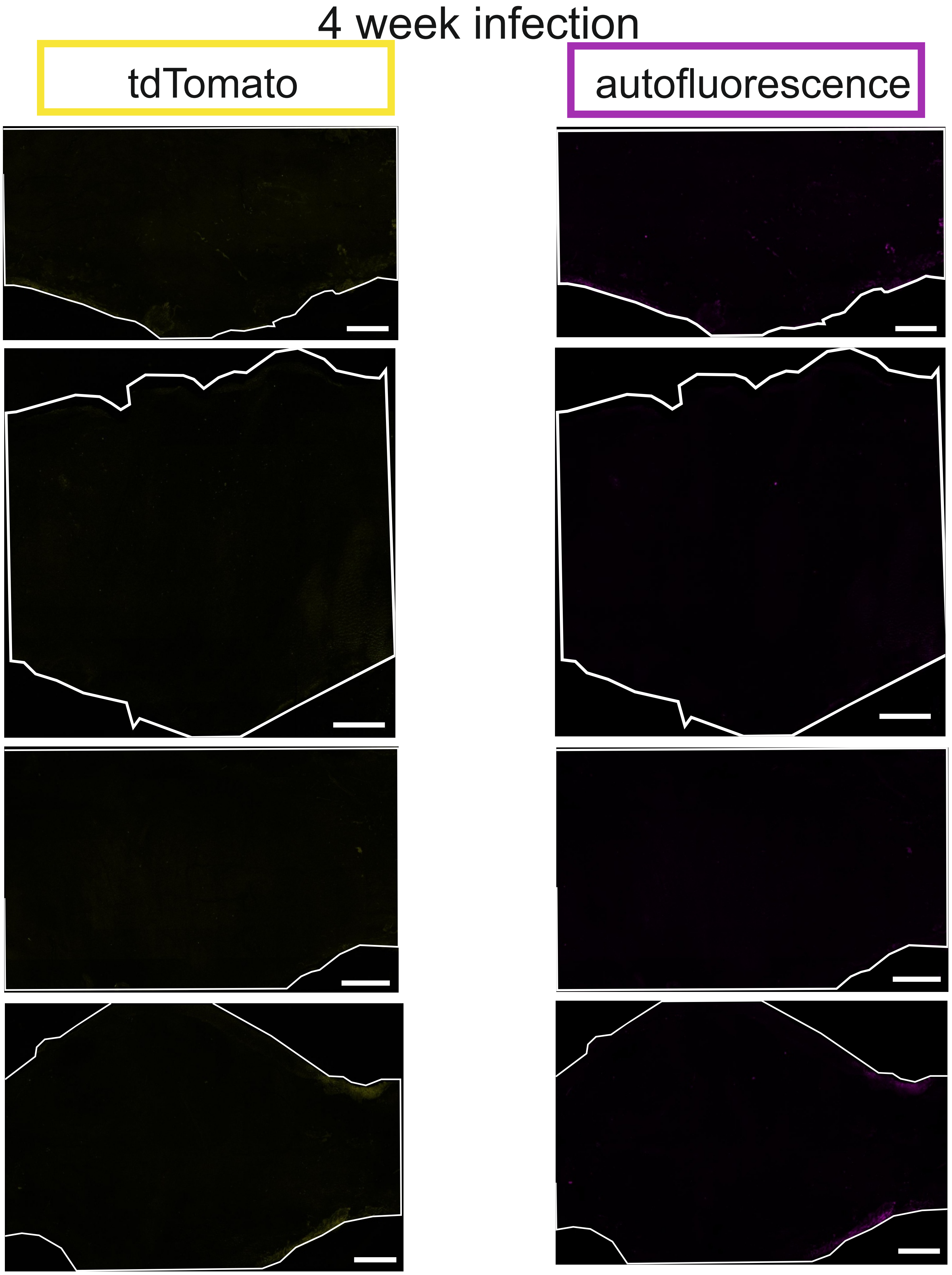


**Supplementary Figure 1: Detection of tdTomato-expressing *L. donovani* in the skin of C57BL/6J mice at 4 weeks post-infection.** Whole skin images (after stitching of panels) of infected skin at 4 weeks post-infection (n=4 female mice). tdTomato signal shown in yellow. Autofluorescence signal shown in magenta. Scale bar represents 5000µm.
