## Supplementary Figure 2 for "Bi-directional highways and super-seeder tissues underpin parasite dissemination in experimental visceral leishmaniasis"

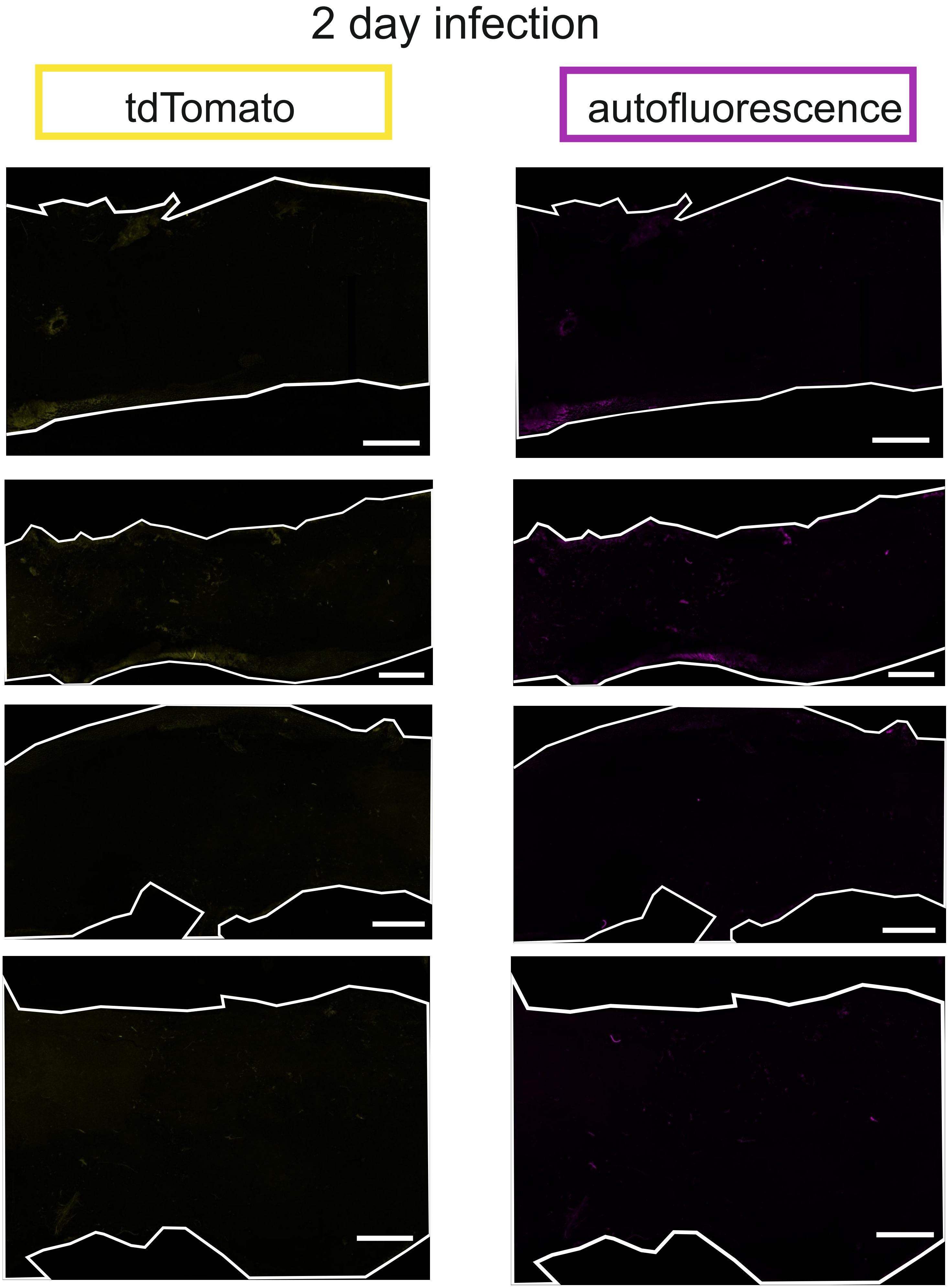


**Supplementary Figure 2: Detection of tdTomato-expressing *L. donovani* in the skin of C57BL/6J mice at 2 days post-infection.** Whole skin images (after stitching of panels) of infected skin at 2 days post-infection (n=4 female mice).
