## Supplementary Figure 3 for "Bi-directional highways and super-seeder tissues underpin parasite dissemination in experimental visceral leishmaniasis"

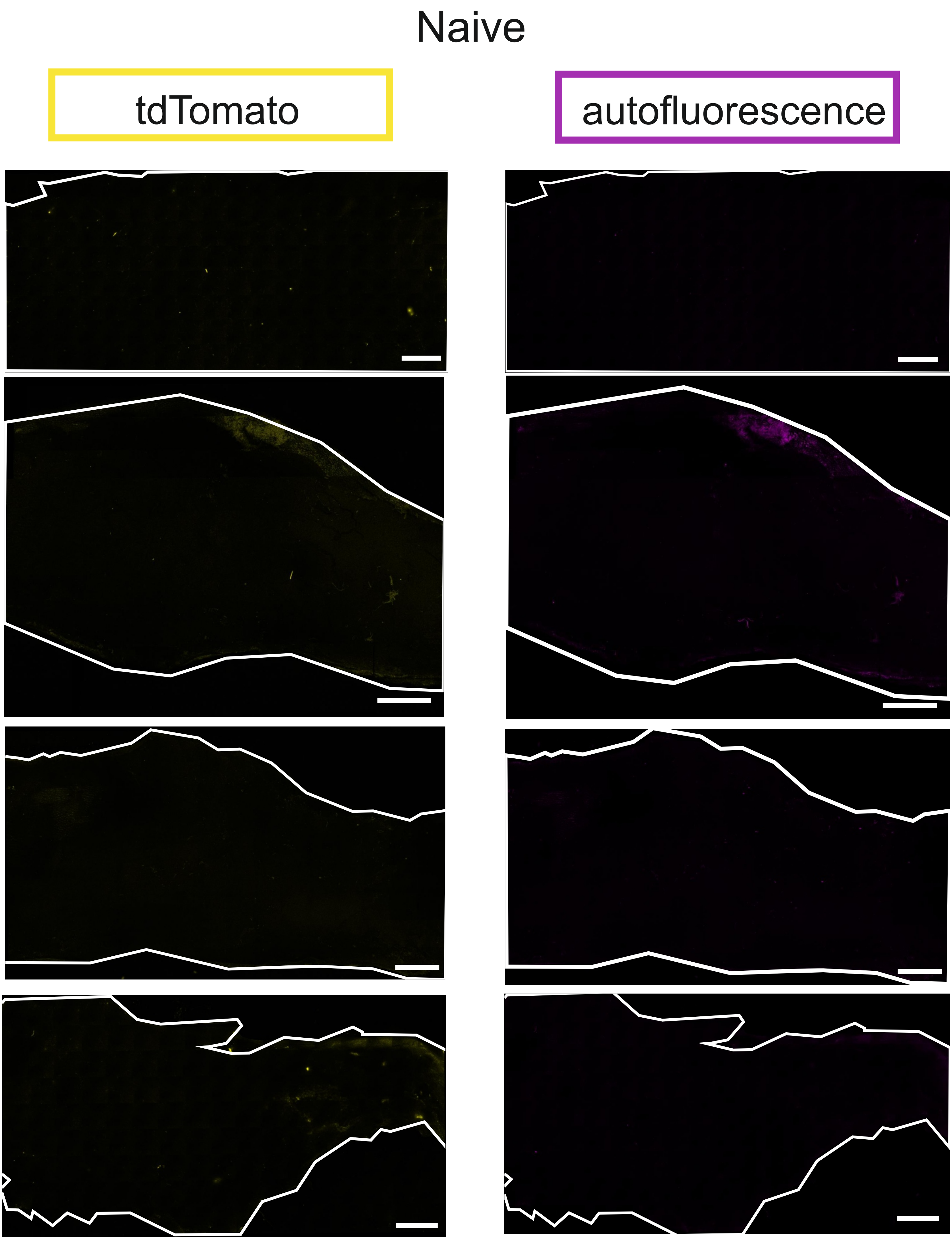


**Supplementary Figure 3: Detection of autofluorescence signal in the skin of naïve C57BL/6J mice.** Whole skin images (after stitching of panels) of naïve mouse skin. tdTomato signal spectrum background shown in yellow. Autofluorescence signal
