## Supplementary Figure 4 for "Bi-directional highways and super-seeder tissues underpin parasite dissemination in experimental visceral leishmaniasis"

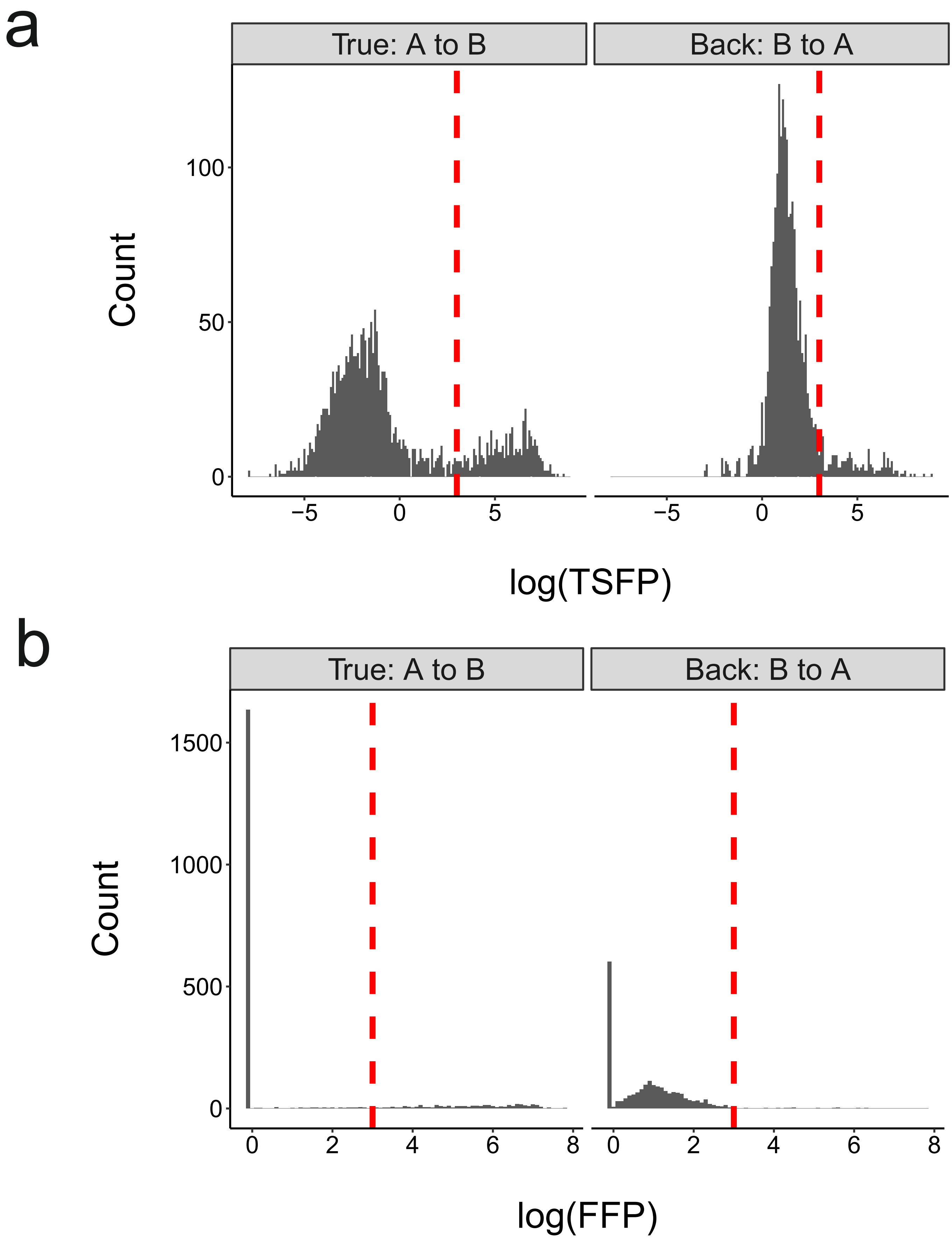


**Supplementary Figure 4: Thresholding histograms for true and back migration between INPUT and samples. a, b,** Histograms for TSFP (a) and FFP (b) for true migration (INPUT samples as population A, tissue samples as population B) and back migration (tissue samples as population A, INPUT samples as population B). Red dotted line indicates the threshold applied, based on distribution shifts. All FFP below 1 were changed to 0.9 for visualisation. All TSFP of negative value or NA were changed to 0 for visualisation.
