## Supplementary Figure 5 for "Bi-directional highways and super-seeder tissues underpin parasite dissemination in experimental visceral leishmaniasis"

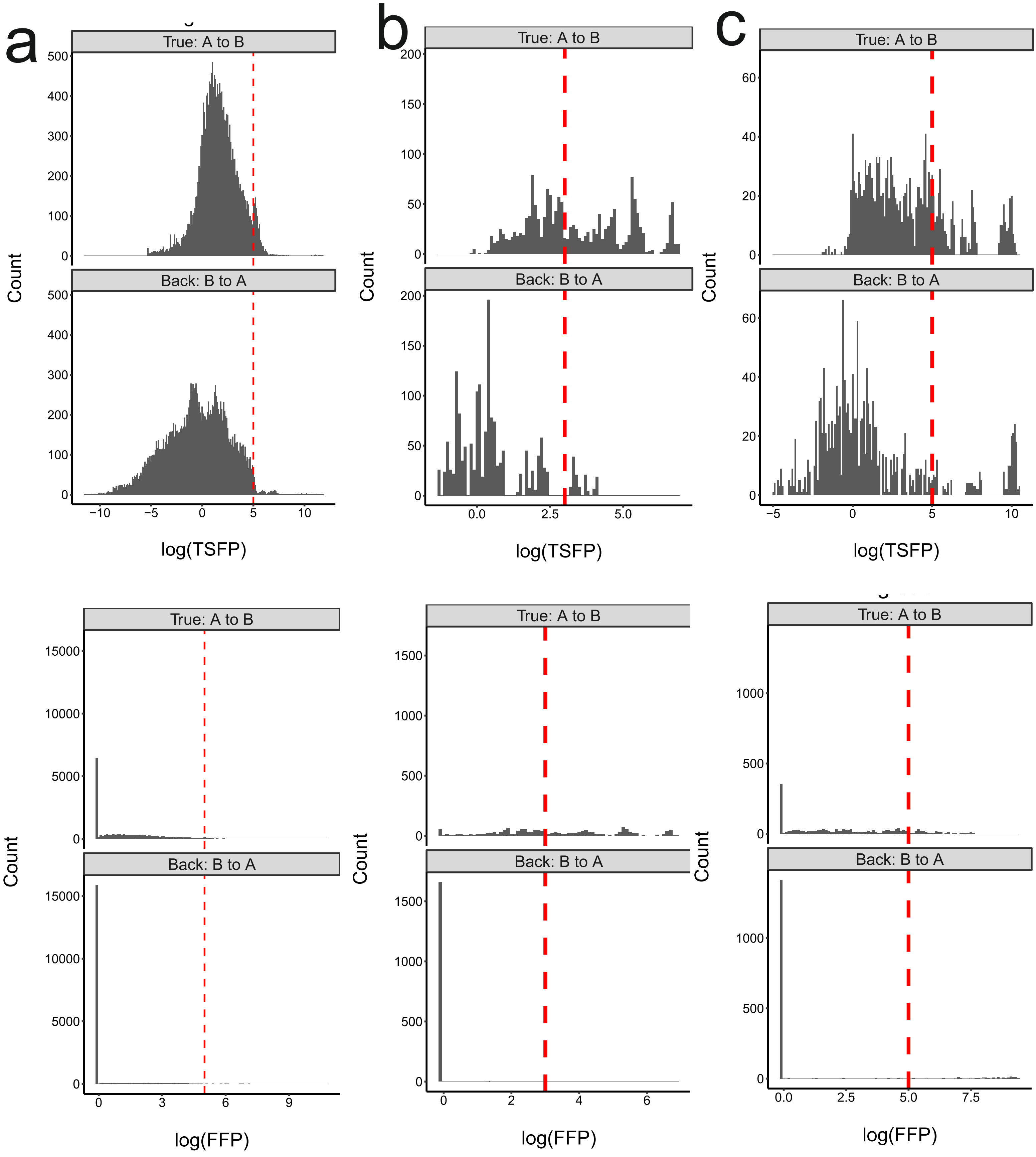


**Supplementary Figure 5: Application of TSFP and FFP to three published datasets. a, b, c,** Histograms for TSFP and FFP for true migration (INPUT samples as population A, tissue samples as population B) and back migration (tissue samples as population A, INPUT samples as population B). Data taken from Hotinger et al (a), Lebrun-Corbin et al (b) and Hullahalli and Waldor (c). Red dotted line indicates the threshold applied, based on distribution shifts. All FFP below 1 were changed to 0.9 for visualisation. All TSFP of negative value or NA were changed to 0 for visualisation.
