## Supplementary Figure 6 for "Bi-directional highways and super-seeder tissues underpin parasite dissemination in experimental visceral leishmaniasis"

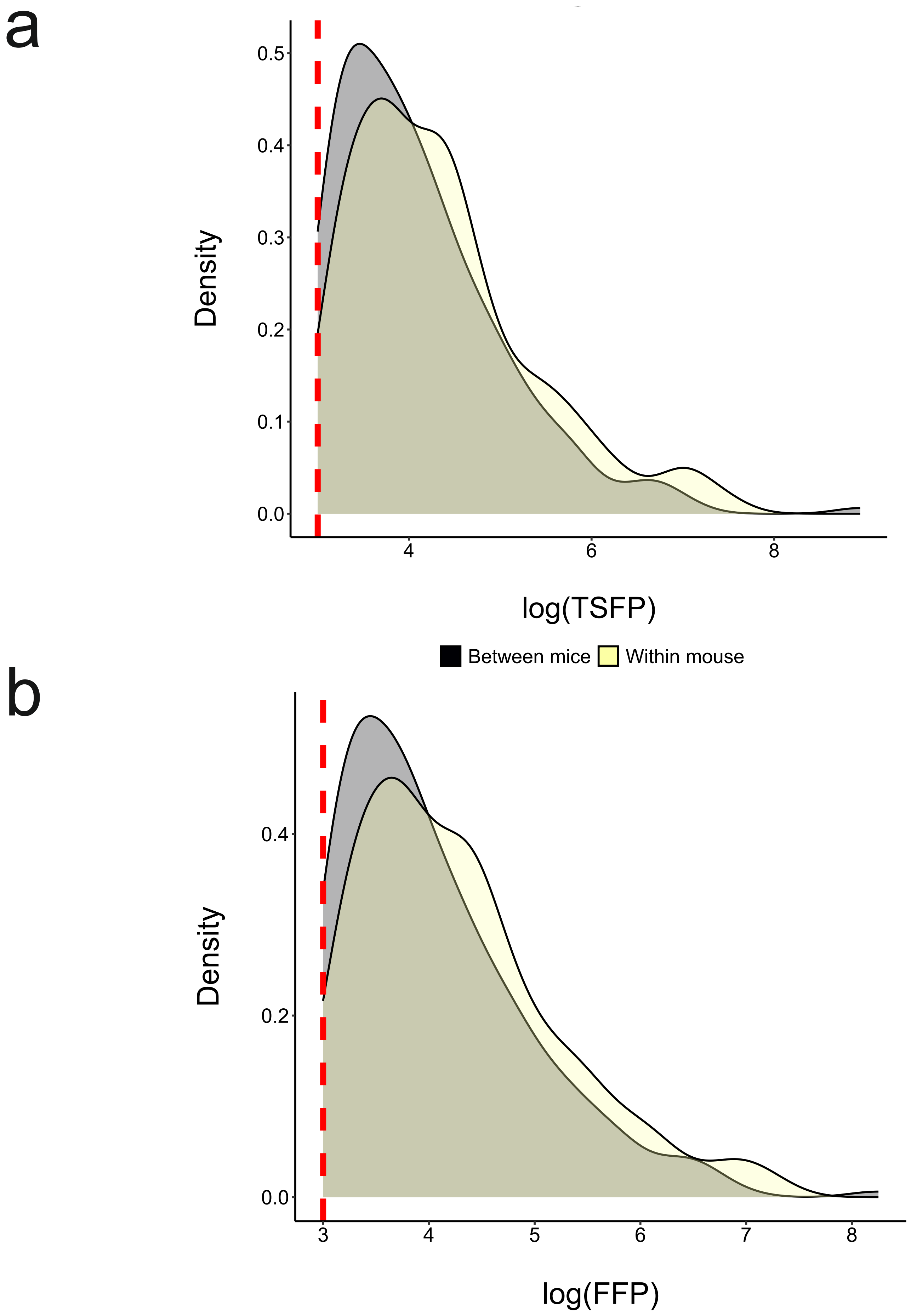


**Supplementary Figure 6: Comparison of within-mouse vs randomised between mouse tissue samples.** a, b, Density plots of TSFP (a) and FFP (b) distributions for all within-mouse (yellow) and a randomly selected matched number of between mouse (grey) tissue samples. Values above the cut-off threshold of 20 are shown.
